# The path to dependence: stepwise genome evolution from a facultative symbiont to an endosymbiont in the N_2_-fixing diatom-*Richelia* symbioses

**DOI:** 10.1101/2025.03.25.645212

**Authors:** V. Grujcic, M. Mehrshad, T. Vigil-Stenman, D. Lundin, R.A. Foster

## Abstract

A few genera of diatoms are widespread in the oceans and form stable partnerships with N_2_-fixing filamentous cyanobacteria *Richelia* spp. A unique feature of the diatom-*Richelia* symbioses is the symbiont cellular location spans a continuum of integration (epibiont, periplasmic, endobiont) that is reflected in the symbiont genome size and content. In this study we analyzed genomes derived from cultures and environmental metagenome-assembled genomes focusing on characters indicative of genome evolution. Our results show an enrichment of short length transposable elements (TEs) and pseudogenes in the periplasmic endosymbiont genomes, suggesting an active and transitionary period in genome evolution. In contrast, genomes of endobionts exhibited fewer TEs and pseudogenes, reflecting advanced stages of genome reduction and increased host dependency in the endobionts. Pangenome analyses identified that endobionts streamline their genomes and conserve a majority of their genes in the core, whereas periplasmic endobionts and epibionts maintain larger flexible genomes, suggesting increased genomic plasticity. Functional gene comparisons with other N_2_-fixing cyanobacteria revealed that *Richelia* endobionts have similar patterns of metabolic loss but are distinguished by absence of specific pathways (e.g. cytochrome bd ubiquinol oxidase, lipid-A) that increase dependency and direct interaction with their respective hosts. In conclusion, our findings underscore the dynamic nature of genome reduction in N_2_-fixing cyanobacterial symbionts and demonstrate the diatom-*Richelia* symbioses as a valuable and rare model to study genome evolution in the transitional stages from a free-living facultative symbiont to a host-dependent endobiont.

## Introduction

Some of the most striking symbioses occur widespread in the surface ocean and involve diverse single celled eukaryotes (protists) which host a broad group of symbiotic partners, including bacteria, archaea, and other eukaryotes.^1–3^ Despite their ecological importance, the specificity, functional roles, and evolutionary trajectories for many of these planktonic symbioses remain poorly understood. Few systems can be maintained in stable culture for extended periods (>2 years), and this limitation has hindered the establishment of robust model systems for studying planktonic symbioses. While more than a century of observations has highlighted the prevalence of planktonic symbioses and their importance in nutrient cycling^4,5^, detailed genomic and evolutionary studies on protist symbioses remain sparse. Yet, the three known examples of organellogenesis: mitochondria, chloroplast and the recently discovered nitroplast, or N_2_ fixing organelle, are derived from endosymbiotic events involving protists.^6–8^ Thus, modern planktonic symbioses provide unique opportunities for studying genome and organelle evolution.

One fascinating microbial protistan symbiotic system involves a few genera of diatoms as hosts and three species of the heterocyst-forming cyanobacteria *Richelia*, as symbionts.^9^ Heterocysts are specialized cells for N_2_ fixation, thus the role of *Richelia* as a nitrogen source is obvious and has been shown on the cellular level.^10^ Diatoms are widespread and highly diverse photosynthetic eukaryotic microalgae that are important primary producers in the modern oceans and contribute to carbon burial.^11^ Genomic and ultrastructural evidences suggest diatom chloroplasts are derived from a secondary endosymbiosis involving a red or green algal endosymbiont.^12^ Furthermore the extensive diversification of diatoms is attributed to horizontal gene transfer (HGT) events, which have incorporated a high number of bacteria-derived genes, and additionally, diatom-specific transposable elements (TEs) that have facilitated both gene acquisitions and losses.^13^

The sequencing of the first four draft genomes of the symbiotic *Richelia* spp. highlighted how the symbiont’s cellular location has influenced its genome size and genetic content.^14,15^ The two endosymbiotic *R. euintracellularis* strains possess the smallest genomes (ReuHH01; 3.2 Mbp, ReuHM01; 2.2 Mbp) with the lowest guanine plus cytosine (GC) content (33.7 and 33.8%, respectively), and lack several N-assimilatory pathways common to free-living heterocystous cyanobacteria.^14^ The partially integrated endosymbiont, *R. intracellularis,* that resides in the periplasmic space of its host diatom has a slightly larger genome and higher GC (RintRC01; 5.48 Mbp; 39.2%). *Richelia rhizosoleniae* attaches to the outside of the host diatoms (epibiont) and possesses the largest genome with the highest GC (RrhiSC01; 5.97 Mbp and 39.5%); *R rhizosoleniae* is also the only symbiont that can be maintained freely in culture^16^. Several environmental metagenome-assembled genomes (eMAGs) have been reported that are closely related to *Richelia*.^9^

Small genomes are common among obligate symbionts, as are loss of DNA-repair genes and decreased GC content.^17,18^ In the best studied symbiotic systems, symbiont genomes tend to retain essential genes (e.g., genes encoding ribosomal proteins, replication, transcription, translation and cell division), while losing redundant genes of their respective hosts.^19,20^ Earlier comparative genomic studies on symbiotic bacteria of insects and some ciliates^21–23^ identified that the initial stages of genome degradation involve the proliferation of TEs, followed by gene inactivation, pseudogene accumulation, and genomic deletions.^24,25^ TEs can drive genome evolution by enabling the movement and/or interruption, duplication, and rearrangement of genetic information.^26^ As symbionts transition to a more obligate and permanent state (e.g., towards an organelle state) the prevalence of TEs and intergenic regions tend to decrease in their genomes.^20^ Over time, ongoing deletions remove pseudogene fragments and gene loss continues, resulting in small and more compact genomes.^27,28^

The diatom-*Richelia* symbiosis provides an ideal system for investigating the evolutionary trajectory of symbiont genomes due to its continuum of cellular integration. However, the impact of this integration on symbiont genome content remains poorly characterized. In this study, we analyzed four *Richelia* spp. draft genomes and ten eMAGs using comparative genomics and pangenome analyses. We focused on evidence of ongoing genome reduction processes, including changes in intergenic spacer size, the presence of transposable elements (TEs), and pseudogenes. Finally, we compared the functional gene content of *Richelia* with other obligate endosymbiotic cyanobacteria, including unicellular types and symbionts of non-diatom hosts (e.g., spheroid bodies of freshwater diatoms (EtSB), and the hornwort symbiont *Nostoc azollae*) and the UCYN-A/nitroplast, to identify common patterns of functional retention and loss in N₂-fixing symbionts.

## Results and discussion

### Diversity and specificity of Diatom-*Richelia* symbiosis

The four *Richelia* spp. draft genomes (ReuHH01, ReuHM01, RintRC01, RrhiSC01) and ten eMAGs were all placed among the *Richelia* genus in the GTDB phylogeny^29^ and separated according to host (Figure 1). This phylogeny agrees and expands upon earlier studies.^9,31,32^ We used a >95% pairwise average nucleotide identity (ANI) threshold for classifying the various eMAGs according to their expected cellular location (endobiont, periplasmic, epibiont) and associated host. Three additional eMAGs together with epibiont *R. rhizosoleniae* (hereafter RrhiSC01) form a sister clade to the periplasmic and true endobionts (Figure 1). However, only RrhiSC01 is considered as an epibiont as the other three eMAGs cluster but do not have a high ANI and are considered free-living (Figure 1). The five eMAGs that form a clade with the two known *Richelia* endobiont genomes (ReuHH01 and ReuHM01) possess >95% ANI and are considered to be endobionts. The DT104 MAG that clustered with RintRC01 at 98% ANI is considered a periplasmic endobiont. One additional eMAG (TARA_PON; *Candidatus* Richelia exalis), reported from the *Tara* Oceans project^33^ formed a sister clade to the endobionts, however, ANI was < 95%, and therefore the symbiont cellular location and host association remain unknown (Figure 1, Supplementary table 1). We do however, include TARA_PON and the three free-living eMAGs in the analyses for comparison with their closest *Richelia* spp. relative (RintRC01 and RrhiSC01, respectively).

**Figure 1.**
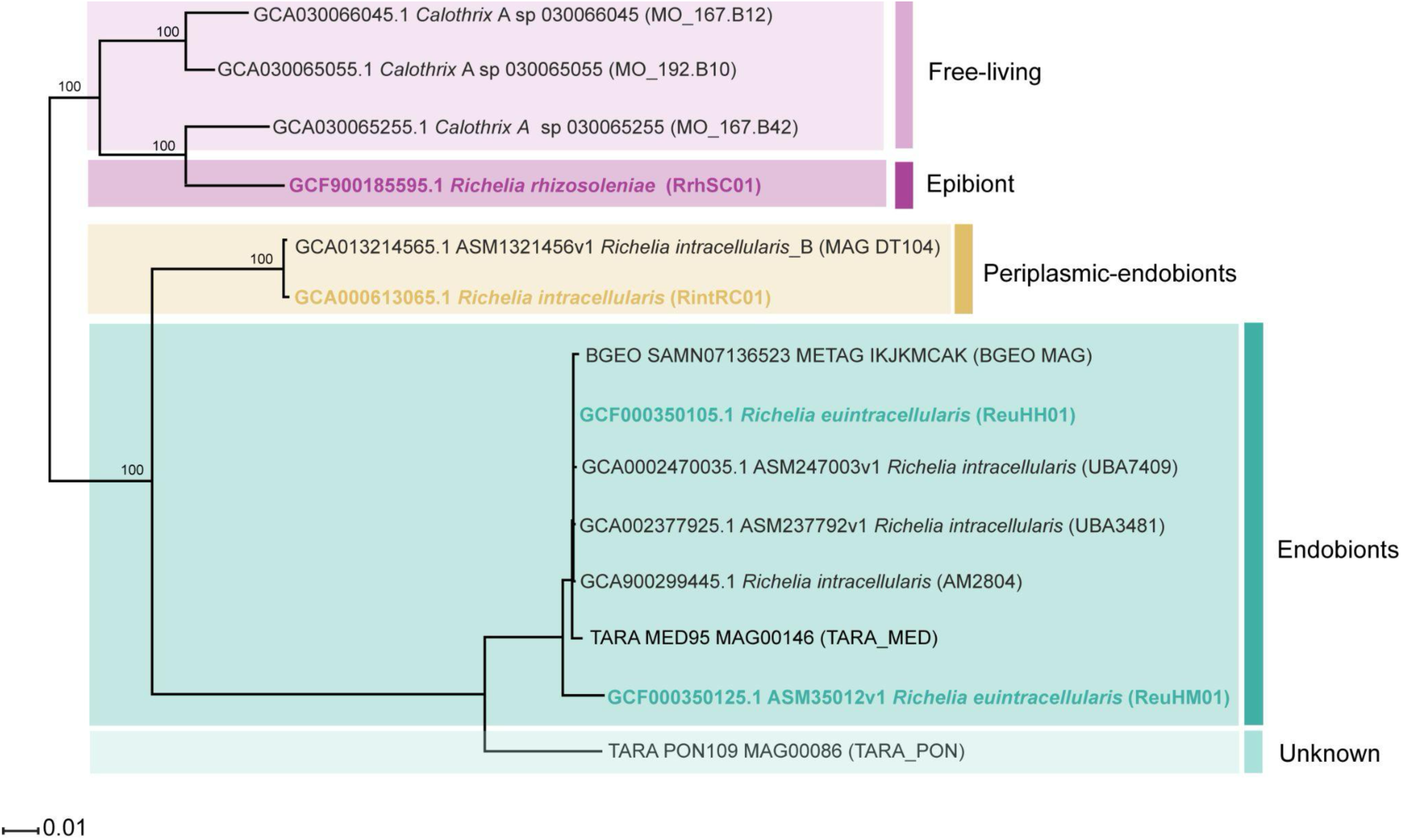
Phylogeny of *Richelia* spp. and its closest relative genera *Calothrix_A.* The draft reference *Richelia* spp. genomes derived from cultured isolates are highlighted in bold text and eMAGs are named according to the GTDB database. Shortened identifiers used throughout this study are shown in parenthesis after the genome names. MAGs are colored according to their presumed cellular location with their respective host diatoms: free-living (light purple), epibiont/facultative symbiont (purple), periplasmic or partial endobiont (yellow), endobionts (green) and unknown (light green). The phylogeny used GTDB-Tk^30^ to place the genomes and eMAGs in the GTDB taxonomy (Parks et al. 2018)^29^.

### Coding and noncoding fractions reflect genome degradation stages in *Richelia* symbionts

Genome statistics were calculated for the *Richelia* draft genomes and eMAGs (Figure 2, Supplementary Table 1). There is a direct correlation between genome size and GC content in the *Richelia* spp.: endobionts possess smaller genomes and lower GC content (3.39Mb ± 0.34 and 34.03% ± 0.88) compared to periplasmic endobionts (5.17Mb ± 0.78 and 39% ± 0.07) and the epibiont (5.98 Mb and 40%) (Figure 2a, Supplementary table 1). The number and percentage of coding sequences (CDSs) follows a similar trend where genomes of endobionts have fewer CDSs (2038 ± 175; 56% ± 4.81) compared to periplasmic endobionts (6029 ± 1548, 67% ± 1.50) and the epibiont (4954; 76.09%) (Figure 2b-c, Supplementary Table 1, Supplementary figure 1a).

**Figure 2.**
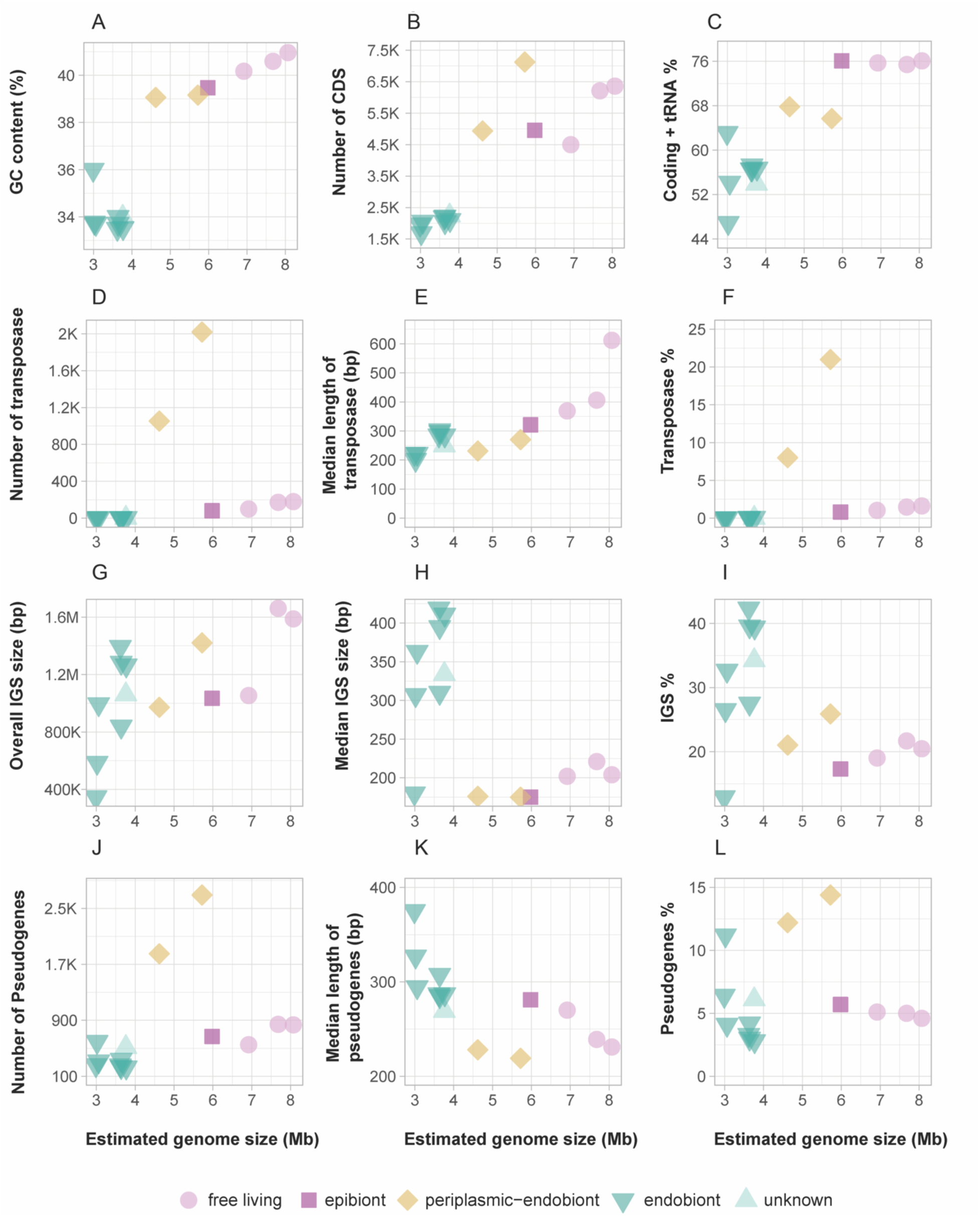
Summary of genomic features for the reference *Richelia* spp. genomes and eMAGs in relation to genome size (Mbp). Each panel (a-l) represents a different genomic character plotted against genome size: a) GC content (%), b) number of coding sequences (CDS), c) overall CDS + tRNA of the genome (%), d) number of transposases, e) median length of transposases (bp), f) overall length of transposases as a percentage (%) of the genome, g) overall length of intergenic spacers (IGS) (bp), h) median length of IGS (bp), i) overall length of IGS as a percentage of the genome, j) number of pseudogenes, k) median length of pseudogenes (bp), and l) overall length of pseudogenes as a percentage (%) of the genome. Genomes are grouped according to their cellular location: free living (circles), epibiont (square), periplasmic-endobionts (diamonds), endobionts (triangle facing up) and unknown (triangle facing down). Detailed statistics for each genome are provided in Supplementary table 1.

To identify other features indicative of genome degradation, we examined the presence, abundance, and length of TEs. TEs are reported in a majority of bacterial and archaeal genomes^35–37^ where the proportion of TEs in prokaryotic genomes is usually below 3%.^38–39^ Furthermore, TEs tend to proliferate in endosymbiotic microbial genomes, particularly those that have recently transitioned to a host-restricted lifestyle.^39^ The number (1.56 ± 1.13), median length (231 ± 107 bp), and proportion of TEs in the genomes (0.02 ± 0.02%) of the endobionts (Figure 2d-f, Supplementary table 1) were very low and similar to the TE content reported for other obligate symbionts (e.g., symbionts of insects, clams, and amoebae).^40–42^ The TE characteristics for the epibiont were similarly low (Figure 2d-f, Supplementary table 1). The periplasmic endobiont genomes, however, contained the highest number and median length of TEs (1537 ± 684 and TEs and 251 ± 28 bp, respectively) and resulted in an unusually high percentage of their genome (14.58 ± 9%) (Figure 2d-f, Supplementary table 1; Supplementary figure 1c). Given these latter results, we further compared the TEs of the periplasmic endobionts to those of other microbial endosymbionts reported with high TE genome content (e.g. *Wolbachia*; Supplementary table 2). Notably, the one periplasmic genome (RintRC01) contained the highest detected percentage (21%) of TEs of any known prokaryote. The periplasmic endobionts also showed an unusual profile for TE length (Supplementary figure 2). Although TE-rich genomes typically contain a high number of full-length TEs, those genes were relatively rare in the two periplasmic endobionts (RintRC01 and MAG DT-104), e.g., 65 and 17 instances, respectively. Instead, the majority of the TEs were short fragments and most TEs were less than 15% of the full length. Upon closer inspection, the fragments were also in clusters (Supplementary figure 2a-b), suggesting they are the remains of highly degraded TEs or the result of multiple TE insertions. This pattern is similar to what has been observed in other endosymbionts, such as the *Wolbachia* bacteria pathogen, where TE-rich genomes accumulate numerous truncated and degraded TEs, likely due to reduced transpositional activity after an initial phase of proliferation.^43^ The low numbers of full length and potentially active TEs compared to the high numbers of fragments suggest that the TE proliferation in the periplasmic endobionts has started to decline after a period of high activity.

Following examination of CDSs, we calculated statistics for intergenic spacers (IGS), which tend to be less constrained by selective pressure compared to CDS. The IGS length followed the same trend as the other genome statistics: longest in the epibiont (1,034,196 bp), followed by periplasmic endobionts (1,196,254 ± 317,383 bp) and endobionts (957,936 ± 310,050 bp) (Figure 2g, Supplementary table 1). However, according to the median size of individual IGSs, the endobionts (341 ± 84 bp) were distinguished from the other symbionts (Figure 2h, Supplementary table 1). Furthermore, the IGS comprised a significant fraction of the endobiont genomes and accounted for 31.55% ± 10.26 of the assembly length (Figure 2i, Supplementary table 1), which is approximately twice the percentage commonly reported for free-living bacteria^44^ and higher than the IGS portion of the endobiont and the epibiont genome assemblies (23.46% ± 3.42 and 17.31%, respectively) (Figure 2i, Supplementary Table 1). However, it is important to note that the overall IGS proportion varied greatly among endobiont genomes (12-42% (Figure 2i), and suggests different stages of genome degradation among the endobionts, which might be expected as the MAGs are derived from environment. Genomes with the least IGS could already be in advanced stages of genome degradation with an increased host dependency.

Variation of the IGS in the *Richelia* genomes raises important questions about the role of pseudogenes in their genome degradation. Pseudogenization is a key mechanism for gene loss, and results from the accumulation of mutations in protein-coding sequences and can often lead to introduction of premature stop codons.^17,45^ In prokaryotes, pseudogenes typically make up between 1-5% of the genome^46^, indicating there is purifying selection to keep genes functional.^45^ However, intracellular pathogens and endosymbionts in transitional stages of genome reduction often exhibit high numbers of pseudogenes (10 - 50% of their genome) which reduce their coding capacity significantly.^47,48^

The prevalence of pseudogenes varied in the *Richelia* genomes and eMAGs (Supplementary table 3). As expected, the highest number of pseudogenes was found in the partial endobionts (2,272 ± 594), followed by the epibiont (669) and endobionts (318 ± 128) (Figure 2j, Supplementary Table 1). While endobionts contained fewer pseudogenes overall, they had the largest median pseudogene size (309 ± 33 bp) (Figure 2k, Supplementary Table 1), likely due to a slower rate of DNA loss, a phenomenon documented in other endosymbiotic bacteria. For instance, in *Buchnera aphidicola*, an obligate endosymbiont of aphids, pseudogenes can persist for extended periods, with a mean half-life estimated at 24 million years, indicating a gradual process of genome reduction^.48^ Despite their smaller pseudogene size, periplasmic endobionts had the highest proportion of pseudogenes relative to genome assembly size (13.3 ± 1.5%) which was a significantly larger fraction compared to the epibiont (5.75%) and endobionts (5.02 ± 2.97%) (Figure 2l, Supplementary Table 1). We interpret this high prevalence of pseudogenes in the periplasmic endobiont genomes as indicative of relaxed selection on many genes and functions coupled to either a shorter time since pseudogenization or weaker selection for genome streamlining. In contrast, the low number of pseudogenes and fewer CDSs in the endobiont genomes suggests stronger selection for genome streamlining, possibly to minimize replication costs during cell division and decreased redundancy with functions in the host diatom. Additionally, over 40% of IGS in some of the endobionts were composed of detectable pseudogenes (Supplementary Table 1, Supplementary Figures 1e-f, Supplementary figure 3), potentially resulting from a slower rate of deletion or an accumulation of larger, non-functional genomic regions.^49^ The lack of purifying selection for function in IGS regions has led to an increased mutational bias towards AT richness in e.g., endosymbionts of insects and some obligate pathogenic bacteria (e.g., *Rickettsiales* and *Chlamydiales*).^19^ This same pattern in AT richness was also observed in the *Richelia* endobiont genomes (Figure 3).

**Figure 3.**
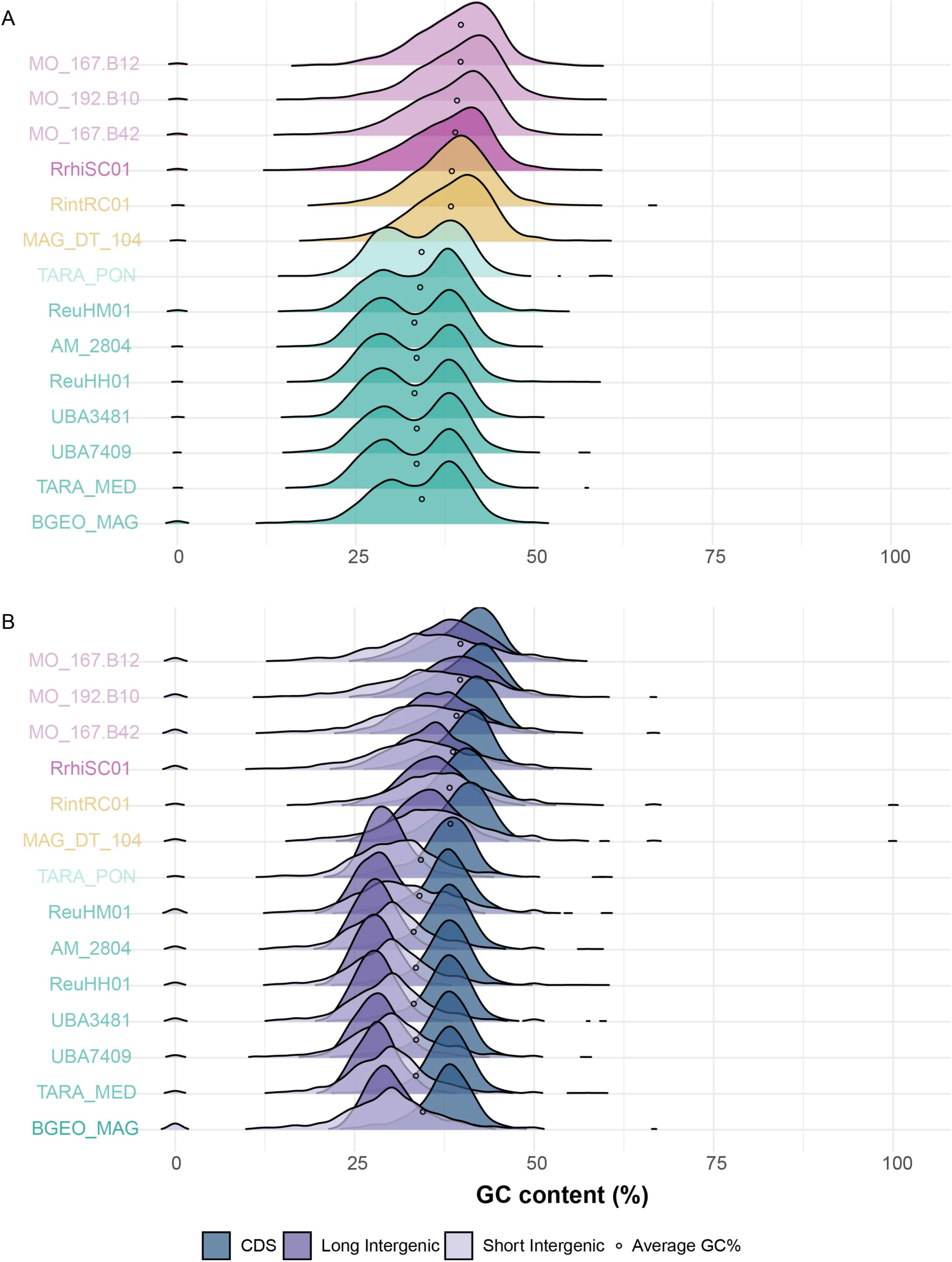
Distribution of GC content in coding sequences and intergenic spacers in *Richelia* spp. genomes and eMAGs. a) Overall GC content distribution for each genome represented by a density plot as a percentage (%). Genomes are color-coded according to their cellular location: free-living (light purple), epibiont/facultative symbiont (purple), periplasmic or partial endobiont (yellow), endobionts (green) and unknown (light green). b) GC content was calculated separately for coding sequences (CDS), long intergenic regions (<300 bp), and short intergenic regions (>300 bp). Each genomic category is represented by a density plot, with different colors indicating the type of genomic feature: CDS (blue), long intergenic regions (purple), and short intergenic regions (light purple). The average overall average GC content for each genome is indicated by a black dot.

### Lower GC content in endobionts is driven by non-coding regions

The distribution frequency of GC content in a genome can reveal underlying processes such as mutational biases and selection pressures.^50^ Our analyses showed a unimodal distribution of GC content in *Richelia* epibionts and periplasmic endobionts. The *Richelia* endobionts, however, exhibit a bimodal distribution (Figure 3a). By further separating CDSs and IGS and categorizing IGS by length (with a 300 bp threshold), we identified in the endobionts that the higher GC peak corresponds to CDS and the lower GC peak is primarily found in the IGS (Figure 3b, Supplementary Figure 4). This bimodal pattern, associated with genome reduction in obligate symbionts, reflects increased genetic drift and relaxed selection pressures.^17,20^ The median sizes of IGS (as described in above) in endobionts are longer compared to partial endobionts and epibiont, which, alongside fewer CDSs, contributes to their lower overall GC content. In contrast, the median CDS length is 767 bp in the epibiont, 374 ± 66 bp in periplasmic endobionts, and 668 ± 94 bp in endobionts (Supplementary Table 1). The low relative GC content in IGS and their low similarity to known genes are indicative of *Richelia* endobionts being in advanced stages of genome reduction, a result common to other symbionts in prolonged relationships with their hosts.^49,51^

### Symbiotic lifestyle shapes the pangenome of *Richelia*

Our pangenome analysis of the *Richelia* genomes and closely related eMAGs identified 11,447 unique gene clusters from a total of 52,462 genes (Figure 4). The unique gene clusters were categorized into eight bins based on their occurrences across the different genomes and eMAGs. The core genome bin contained 1,768 gene clusters (15.4%) present in all genomes and eMAGs, and we identified six flexible genome bins based on their distribution in the pangenome (37%): flexible 1 (n=158), flexible 2 (n=668), flexible 3 (n=1,041), flexible 4 (n=639), flexible 5 (n=507), and flexible 6 (n=1,212) (Figure 4). Finally, gene clusters unique to individual genomes were categorized as singletons (n=5,454; 47.6%) (Figure 4).

**Figure 4.**
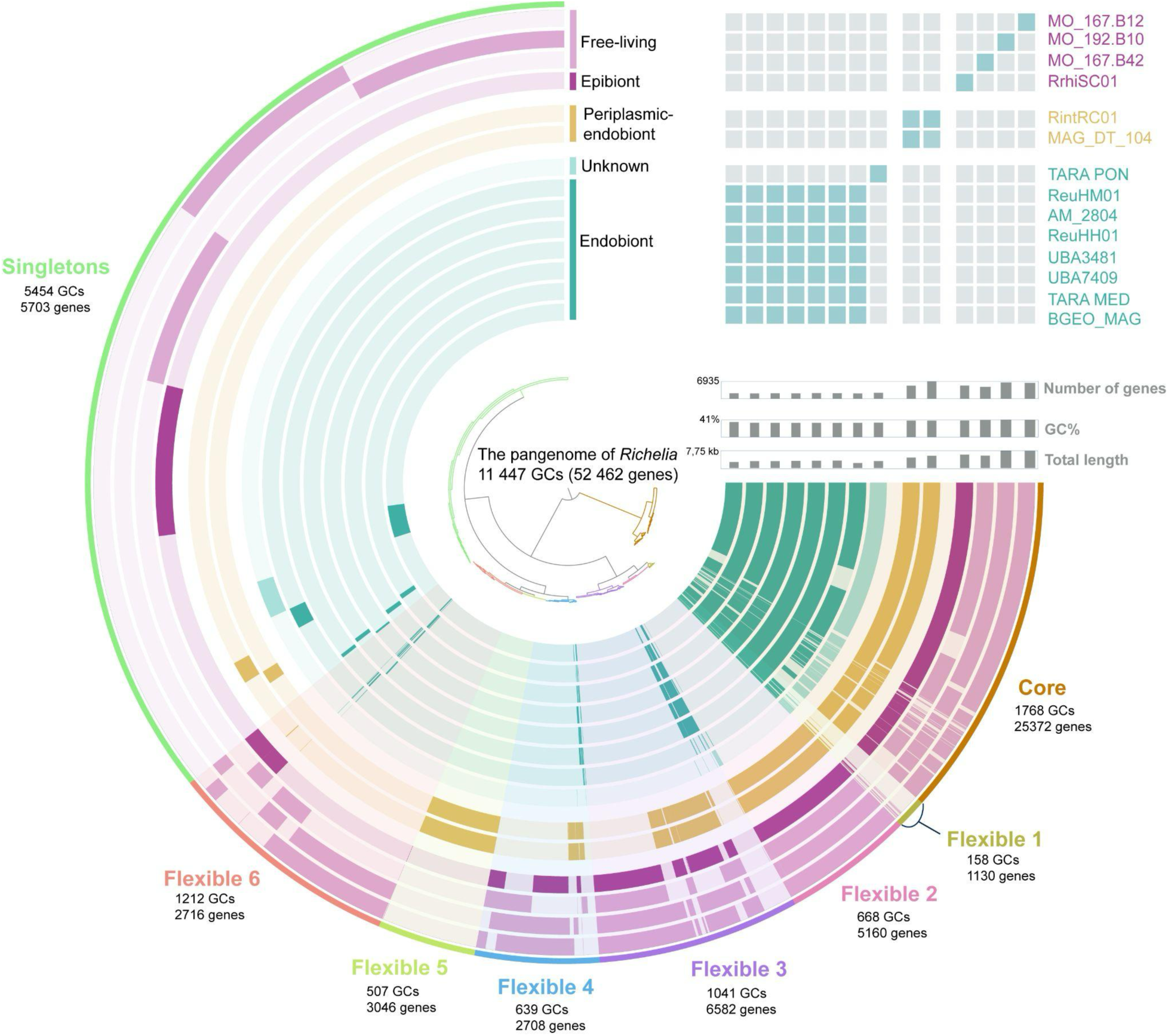
The pangenome of genus *Richelia* spp. and close relatives. Pangenome covers 52 462 genes and 11 447 gene clusters from fourteen genomes of *Richelia* and eMAGs corresponding to five different cellular locations: free living, epibionts, periplasmic endobionts, endobionts and unknown. Top right corner illustrates the average nucleotide identity (95%) that groups genomes into eight species. Genomes were organized based on their placement in the phylogenetic tree illustrated in Figure 1.

The core genome represents functions conserved across all *Richelia* strains which are critical for their survival.^52^ The flexible genome likely reflects the adaptability to various environmental conditions, including specifically for *Richelia* their cellular location. The high proportion of singletons (nearly 50%) indicates significant genomic plasticity, suggesting that individual genomes may retain and/or acquire unique functions for specific purposes.^53^ Interestingly, a recent study^54^ highlighted that a bacteria’s lifestyle, particularly the degree of host integration, plays a significant role in shaping pangenome fluidity (also called genome fluidity). Pangenome fluidity refers to the average proportion of genes unique to any two genomes of the same species.^54^ Collectively and expected given the continuum of cellular integration of *Richelia* with its respective hosts, the *Richelia* genomes possessed varying degrees of fluidity. For example, endobionts exhibited a higher proportion of their gene clusters in the core (71.98 ± 4.72%) and far less genes in the flexible (10.85 ± 4.33%) and singleton (6.06 ± 7.54%) parts of the pangenome (Supplementary table 4). Thus, the endobionts possess more conserved and less fluid pangenome structure, which is consistent with their presumably stable, intracellular lifestyle. In contrast, greater pangenome fluidity was observed in the epibiont and periplasmic endobionts, as evidenced by their larger flexible genome fractions (47.32 ± 3% and 49.46 ± 0.32%, respectively) (Supplementary table 4). Unlike the endobionts, both the periplasmic endobionts and the epibiont are adapted to different interactions (e.g., competition, cooperation) with their hosts and a higher instability in their respective cellular environments compared to the intracellular endobionts.

More than 60% of the genes were annotated using the Kyoto Encyclopedia of Genes and Genomes (KEGG) functional categories (Supplementary Figure 5). Most of the annotated gene clusters were part of the core genome, while many of the unannotated genes were associated with the flexible or singleton parts of the pangenome (Supplementary Figure 6). Our comparison of the functional genomic content retained and lost in different parts of the pangenome further underscored the impact of the different cellular locations.

We started by examining functions related to carbon (C) and nitrogen (N) metabolism given that the symbionts function as a N source and N_2_ fixation in heterocystous cyanobacteria is primarily fueled by photosynthesis. We detected the 12 *nif* genes for the nitrogenase complex required for N_2_ fixation in the core genome: *nifH* (K02588), *nifD* (K02586), *nifK* (K02591), *nifE* (K02587), *nifN* (K02592), *nifX* (K02596), *nifB* (K02585), iscS/NFS1 (K04487), *nifT* (K02593), *nifV* (K02594), *nifW* (K02595), and *nifZ* (K02597) (Supplementary figure 7). Out of the 12 *nif* genes, seven genes (*nifUBENXSV*) encode biosynthesis of the iron molybdenum cofactor (FeMo-co) which is essential for the catalytic activity of nitrogenase in the conversion of atmospheric N_2_ to ammonia. One gene, *nifJ* (K03737), which encodes pyruvate (flavodoxin) oxidoreductase (PFOR), was detected only in the flexible 3 of both the periplasmic endobionts and epibiont genomes and the free-living eMAGs but absent in all endobionts genomes. Without *nifJ*, the endobionts may rely solely on the pyruvate dehydrogenase complex (PDC) for pyruvate oxidation under oxic conditions, rather than utilizing PFOR’s flavodoxin-dependent pathway, which is typically important under anaerobic or iron-limited conditions.^55^ In endobionts, *nifJ* gene loss likely reflects an adaptation to the stable environment provided inside the host cytoplasm, where oxygen sensitivity of PFOR and a more stable iron availability could make the ferredoxin pathway sufficient for nitrogenase (or hydrogenase) activity without the need for a flavodoxin-based reduction.

Additionally, genomes of both the periplasmic and true endobionts have lost genes associated with assimilatory nitrate reduction (*narB*, K00367; *nirA*, K00366) that remain in the flexible 3 region of the epibiont RrhiSC01 pangenome and free-living representatives. For comparison, both *narB* and *nirA* are also absent from EtSB and UCYN-A, but *nirA* is retained by *N. azollae* endobionts. These findings highlight the influence of both the host and the persistently oligotrophic natural environment for which these planktonic symbioses, including the diatom-*Richelia* symbioses, persist, on their N assimilation strategies.

Recent evidences have shown that the *Richelia* spp. symbionts, including the periplasmic endobionts, differ in their C metabolic activity and host dependency.^56,57^ To explore this further, we examined the photosynthesis module in KEGG and identified 45 photosynthetic genes in the core, 15 genes in the flexible, and no genes in the singleton part of the pangenome (Supplementary Figure 7). Among the 15 genes in the flexible section, six were not redundant with core genes, and included four genes that encode functions for PSII (*PsbJ*, *PsbP*, *PsbT*, and *Psb28-2*; K02711, K02717, K02718, K08904), one gene (*Psal*, K02696) for PSI, and one gene required for electron transport (*PetJ*, K08906) (Supplementary figure 8). With the exception of *PsbT* (present in flexible1 of some endobionts), all six genes were missing from the genomes of endobionts but present in periplasmic endobionts, the epibiont, and free-living relatives. The notable deletion of *PetJ*, which encodes cytochrome *c6*, in endobionts likely reflects an adaptation to the controlled intracellular environment of the host. In free-living heterocystous cyanobacteria, cytochrome *c6* is essential in heterocysts, where it serves as the primary soluble electron donor to Cox2 under copper-replete conditions—a role that plastocyanin (PetE, K02638, present in all analyzed genomes) cannot fully substitute.^58,59^ In vegetative cells, *PetJ* is typically expressed under copper-limited conditions, when plastocyanin cannot function due to insufficient copper availability.^58^ However, *Richelia* endobionts reside within the diatom host cytoplasm^60^, which could provide both consistent electron donors and sufficient copper that would make *PetJ* redundant. This adaptation allows the endosymbiont to streamline its electron transport system, reducing metabolic redundancy but also increasing its reliance on the host for redox balance.

Another important aspect of photosynthesis is light capture. Light-harvesting proteins of cyanobacteria are organized as antennae called phycobilisomes which are arranged on thylakoid membranes. Interestingly, within the phycobilisome module, 18 of the genes were detected within the core genome and three genes identified in flexible regions of the periplasmic endobiont and epibiont genomes. (Supplementary figure 8). Three of the latter genes were missing in the endobionts and included: *cpcD* (K02287), *cpcE* (K02288) and *cpcF* (K02289), which are involved in synthesis and assembly of phycocyanin, a key component of the phycobilisomes, that captures light energy and transfers it to the reaction centers of the photosystems I and II.^61^ The absence of these genes, along with the loss of some genes in the photosynthesis module (Supplementary figure 8), suggests that endobionts have reduced light-capturing capacity and decreased genome content for their own photosynthetic apparatus. A higher dependency and capacity to transport organic carbon (e.g., sugars) from the host diatom was recently confirmed experimentally and in wild populations of the *Hemiaulus-Richelia* (ReuHH01) symbioses.^56,57^ The latter combined with our functional analyses here provides further evidence for a lowered investment of the endobionts to perform their own photosynthesis.

### Functional comparison with other obligate symbionts highlights gene retention and loss differs amongst N_2_-fixing endobionts

In order to identify how genome reduction and metabolic dependency has evolved in different symbiotic systems involving N_2_-fixing cyanobacteria, we compared the genome content of the *Richelia* symbionts with two unicellular obligate endosymbionts: the spheroid bodies of the freshwater diatom *E. turgida* (EtSB), obligate symbiont that evolved to the organelle UCYN-A/nitroplast, and one obligate heterocystous endosymbiont *N. azollae* of a water fern (Figure 5., Supplementary figure 9).

**Figure 5.**
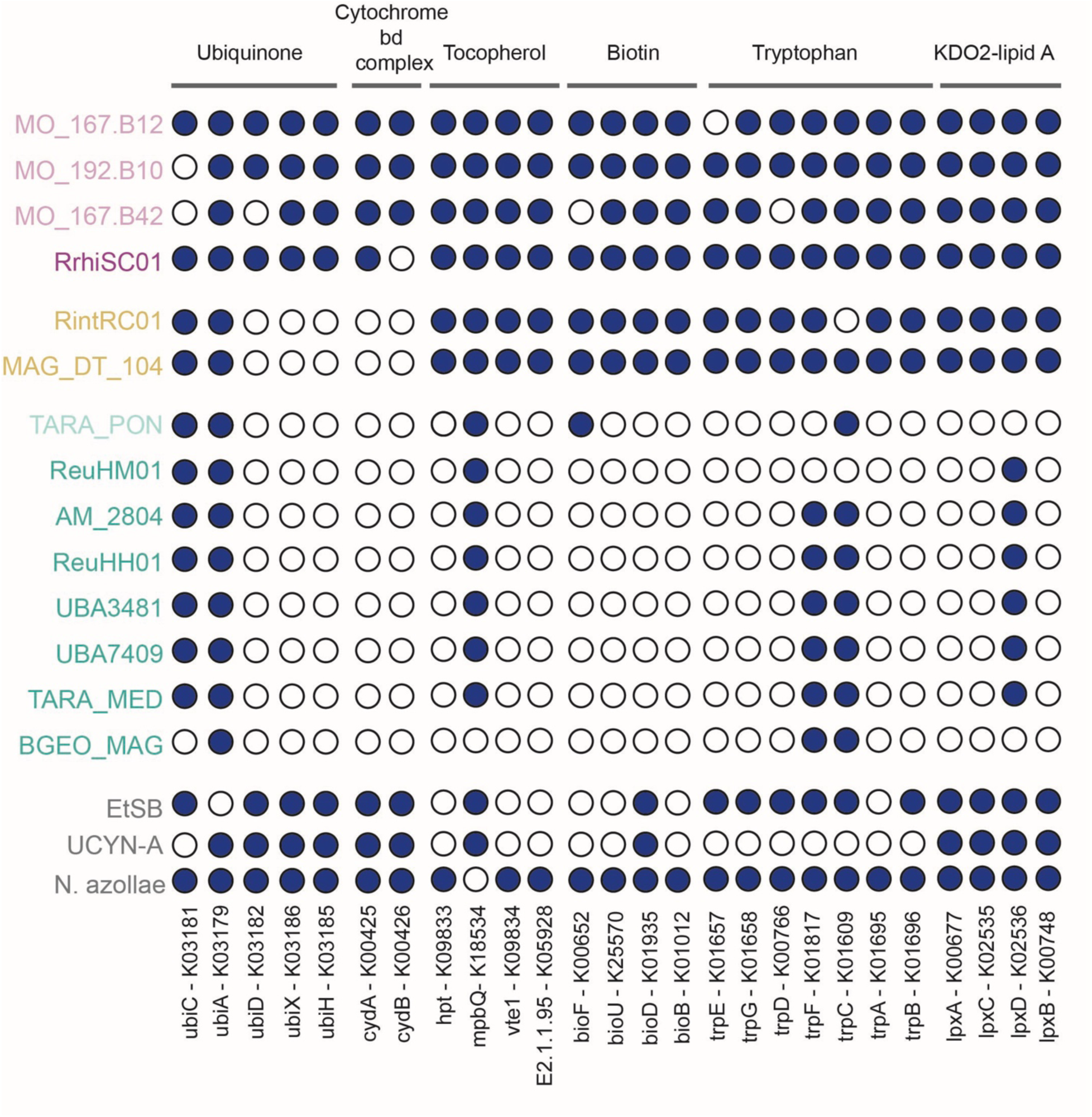
Presence and absence of genes encoding in various biosynthetic pathways. K0 number identifiers together with corresponding gene names are shown as column names. *Richelia* genomes are color coded according to their cellular locations and the three other genomes belonging to obligate N_2_ fixing endobionts and nitroplast (UCYN-A) are colored in gray. Gene presence is marked as a blue circle and absence as a white circle.

Under symbiotic conditions, coding regions that provide little or no added value in a given environment may be lost.^39,62^ Similarly, as observed in bacterial endosymbionts of insects^17,39,62^, repeated population bottlenecks may weaken selection pressures, even for essential genes^19^, and leads to the elimination of dispensable genes through genetic drift.^63^ We identified several such examples, e.g. the *Richelia* endobionts and other cyanobacteria lack several genes for the complete biosynthesis of certain amino acids, vitamins, oxidase in the cytochrome bd complex, and KDO2-lipid A biosynthesis pathway (Figure 5). However, despite their shared N₂-fixing role— and in the case of EtSB, their association with a diatom—the pattern of gene loss and retention was not identical. This variation likely reflects differences in their evolutionary trajectories, host interactions, and genomic constraints, leading to distinct adaptations even within similar symbiotic niches.

The *Richelia* endobionts, including the periplasmic ones, showed several evidences of degraded biosynthetic pathways, including disrupted ubiquinone synthesis and the absence of cytochrome bd ubiquinol oxidase (cytochrome bd complex). Similarly, the genomes of EtSB and UCYN-A/nitroplast also show disrupted ubiquinone synthesis, but retain cytochrome bd (Figure 5). In contrast, both pathways remain intact in *N. azollae*, the epibiont *Richelia* (RrhiSC01), and one of the three free-living eMAGs. Ubiquinone and the cytochrome bd complex are integral components of the aerobic respiratory chain. Thus, the loss of one or both of these components suggests a major disruption in the symbionts’ ability to carry out aerobic respiration independently, limiting their capacity to generate ATP through oxidative phosphorylation, which typically relies on oxygen. We speculate that endobionts are either adapting to low-oxygen environments (not likely given their photosynthetic hosts) or no longer require these functions because they are potentially compensated by the host. In the obligate intracellular *Rickettsia* spp. symbionts, which lack a complete ubiquinone synthesis pathway, the endosymbionts rely on importing compounds from the host to complete several biosynthetic pathways.^64^ A comparable scenario could occur in the various endobionts (*Richelia*, EtSB) and UCYNA/nitroplast, given that one to two of the five *ubi* genes have been conserved for ubiquinone synthesis.

Similar metabolic degradation is seen in the biosynthesis of vitamins, such as α-Tocopherol and biotin. α-Tocopherol is an antioxidant particularly effective at scavenging intracellular singlet oxygen.^65^ The *Richelia* epibiont, periplasmic endobionts and *N. azollae* possess a complete biosynthetic pathway for α-Tocopherol (Figure 5). The *Richelia* endobionts, together with EtSB and the UCYN-A/nitroplast, however, have a degraded pathway with only one gene remaining for α-Tocopherol. We interpret the incomplete α-Tocopherol pathway has resulted from a decreased need for antioxidant defense, and favor a scenario that these endobionts and nitroplast rely on their respective hosts for protection against oxidative stress. For example, the intracellular *Richelia* ReuHH01 is located nearest the mitochondria of their hosts, and in some recent observations possess outer-inner membrane like vesicles in close proximity to the host mitochondria^60^, which suggests a possible mechanism for the host to function in oxidative stress protection.

Biotin is a crucial cofactor for various metabolic enzymes involved in carboxylation reactions, such as fatty acid synthesis, amino acid metabolism, and gluconeogenesis. The biotin biosynthesis pathway in the epibiont RrhiSC01, two of the three free-living eMAGs, and *N. azollae* is complete, containing four genes that encode the necessary enzymes (Figure 5). However, biotin biosynthesis is degraded in the *Richelia* periplasmic endobionts, EtSB, UCYN-A/nitroplast, and completely absent in all *Richelia* endobiont genomes. The inability to synthesize biotin and an intracellular location requires the endobionts to obtain it directly from their host. Most marine algae, including diatoms, can synthesize biotin.^66^ Additionally, several biotin transporters (K03523, K16785, K16786, K16787) were detected in all the *Richelia* symbionts, UCYN-A/nitroplast, and free-living eMAGs (Supplementary figure 10).

We identified that many amino acid synthesis pathways were eroded in the periplasmic and endobiont *Richelia* genomes. Furthermore, we noted that often just one gene was missing, except for tryptophan, where only two out of seven genes necessary for the full pathway remain in the *Richelia* endobiont genomes (Figure 5). Tryptophan synthesis is completely absent in the UCYN-A/nitroplast, and one gene is missing in EtSB, while the full synthesis pathway remains in the other *Richelia* symbionts, their free-living relatives and *N. azollae* (Figure 5). Tryptophan is essential for cyanobacteria in electron transfer and therefore central to capturing sunlight and initiating photosynthesis for efficient energy conversion.^67^ However, tryptophan is the most complex and energy-consuming among all amino acids^68^, which could explain why, this pathway in particular has been extensively degraded in the genomes of endobionts and UCYN-A/nitroplast. It is also likely redundant with their diatom hosts; all diatoms can synthesize tryptophan^69^. Furthermore, amino acids can be imported; *Richelia* symbionts contain homologues of solute binding proteins for N-I and NII amino acid transporters; two of which (e.g., NatB, NatF) have recently been functionally tested and affinities characterized for several substrates.^57^

Finally, the notable loss of genes involved in lipid A biosynthesis in *Richelia* endobionts and retention in the other *Richelia* genomes and their close relatives, EtSB, *N. azollae*, and UCYN-A/nitroplast, suggests a unique loss of structural components in their outer membranes. Cyanobacteria are gram-negative, and possess a cytoplasmic membrane and an outer membrane, which contains lipid A as a key component of lipopolysaccharides.^70^ The pathway that produces lipid A, is uniquely degraded in the *Richelia* endobionts, with only one gene out of four encoding enzymes remaining. The arrangement of the heterocyst envelope of *Richelia* ReuHH01 has also been recently reported as modified^60^ (Figure 5). In contrast, the full lipid A biosynthesis pathway remains in all other genomes, including the other heterocyst-forming *N. azollae*. The loss of lipid A biosynthesis in the *Richelia* endobionts remains unclear, as some endosymbiotic bacteria have lost the capacity for synthesis, while others have not.^71^ In some host-associated bacteria, including several *Gammaproteobacteria*, the loss of the structural barrier in outer cell membranes increases permeability to hydrophobic molecules.^72^ Lipopolysaccharides also stimulate host recognition and immune responses in some well-characterized pathogenic host– microbe interactions of animals and multicellular eukaryotes.^72^ It is unclear if such interactions occur in these planktonic symbioses.

In summary, *Richelia* endobionts exhibit several examples of advanced symbiotic dependency, with genome reduction and loss of many important biosynthetic pathways compared to the periplasmic endobionts, epibiont and free relatives. This places the loss and retention of the functional genome content of *Richelia* endobionts closer to that of the obligate planktonic symbiont EtSB and the UCYN-A/nitroplast, while also still showing distinguishable losses e.g., loss/incomplete cytochrome bd complex, tryptophan, biotin, lipid-A. The functional genome content of the *Richelia* epibiont RrhiSC01 reflects its facultative nature, and the periplasmic *Richelia* appear in a transitory state sharing genome loss/retention with both endobionts and the epibiont strains.

### **Diatom**-*Richelia* symbioses as a model for studying transitionary steps leading to endosymbiosis

While previous studies have reported symbiosis-driven genomic changes, they have primarily focused on symbionts of multicellular organisms such as insects, leeches, worms or plants.^19,73–75^ It remained unclear whether the same trends apply to symbioses between prokaryotes and single-celled eukaryotic protists. The diatom-*Richelia* symbioses are a unique and rare system to study genomic evolution across different cellular integrative stages. For example, the transitionary steps were notable in a stepwise genome degradation and highlighted the crucial role of TEs in reductive evolutionary processes (Figure 6). It remains to be seen whether the *Richelia* will eventually result in the formation of a N_2_-fixing organelle, as is the case of the UCYN-A and the nitroplast.^8^ Further progress in understanding this process will require detailed proteomic studies of the diatom-*Richelia* symbiotic systems. To further elucidate the partner dependency, it will be necessary to sequence the host genomes and experimentally confirm that several of the crucial functions identified as missing are complemented by the host or no longer necessary. In conclusion, we have demonstrated that *Richelia* endobionts have lost many essential functions compared to their periplasmic and epibiotic counterparts, which strengthens the hypotheses that *Richelia* symbionts differ in their dependency and adaptation to their symbiotic life history.

**Figure 6.**
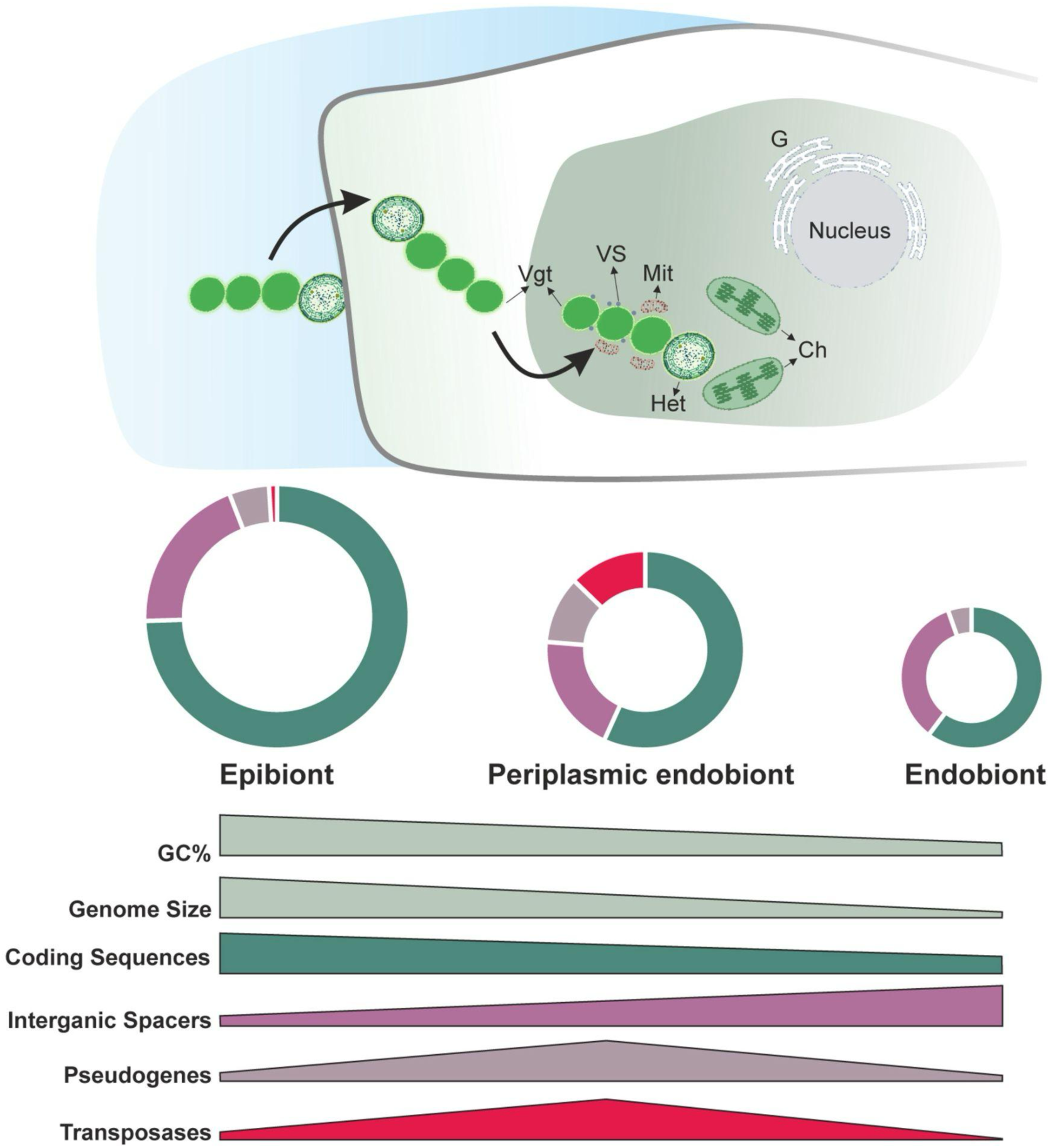
Conceptual view of the evolutionary trajectory of the *Richelia* spp. symbionts according to cellular integration with their respective host diatoms. Upper portion of the figure illustrates *Richelia* filaments, consisting of three vegetative cells (Vgt) and one terminal heterocyst (Het). Epibionts are *Richelia* strains that externally attach to their host diatoms with their heterocysts and have the largest genomes and highest GC%. The periplasmic *Richelia* endobionts reside between the outer cell wall (frustule) and the cell membrane (plasmalemma) of their host diatoms and possess slightly smaller genomes and lower GC%. Finally, *Richelia* are fully integrated into the cytoplasm of their host diatoms and have highly reduced genomes and lowest GC%. Inside of the diatom cell there is a nucleus surrounded by Golgi bodies (G), chloroplasts (Ch) and numerous mitochondria (Mt) in tight associations with *Richelia* endobionts vegetative cells (Vgt). Small dots positioned on *Richelia* endobiont vegetative cells are numerous cell envelope vesicles (Vgt) present in the periplasm which possibly participate in the transfer of metabolites from the cytoplasmic of the cyanobacterium to the diatom. Illustrations of *Richelia* endobionts in the host cytoplasm are based on electron microscopy observations reported recently.^60^ Lower portion of the figure illustrates the genome trajectory of *Richelia* as it transitions from epibionts to endobionts. Epibionts are facultative symbionts and their genomes resemble that of free-living bacteria with high proportion of coding sequence, low proportion of intergenic spacers and few pseudogenes and transposase (as depicted in doughnut plots). Periplasmic endobionts are transitioning from facultative to obligate symbionts and the transition is reflected in their genome characters: slight decrease in the coding sequence and increase in intergenic spacer portions, and substantial increase of pseudogenes and transposases. Genomes of endobionts are characterized by further decreases in the coding sequences and an increase in intergenic spacers. In this stage transposase are nearly purged from the endobiont genomes and there are few pseudogenes accumulated in the intergenic spacers.

## Resource availability

### Lead contact

Requests for further information and resources should be directed toward the lead contact, Vesna Grujcic

### Materials availability

No new materials were generated in this study.

### Data and code availability

Genomes used in this study are all publicly available and their accession numbers are mentioned in the methods and supplementary material. All the materials and code (including supplementary tables 1-4) to reproduce the results of this study are deposited and are publicly available via github: https://github.com/VesnaGr/Richelia_Comparative_Genomics/

## Methods

### Genomes and metagenomes collections

Fourteen genome assemblies were used for this study, 10 of the 14 genome assemblies are reconstructed from environmental metagenomes (i.e., metagenome assembled genome, MAG) (Supplementary Table 1).

### Phylogeny reconstruction

Phylogeny of *Richelia* was extracted from the phylogeny produced by the GTDB-Tk (v. 2.4.0)^30, 76^, which uses pplacer (v1.1.alpha19-0-g807f6f3)^77^ to place genomes in the GTDB taxonomy (R09-RS220).^29^

### Genome statistics

Completeness and contamination of the genome assemblies and MAGs were calculated using CheckM2 (v. 1.0.1).^78^ Genomes were initially annotated with Prokka (v.1.14.6)^79^ including gene recognition and translation initiation site identification with Prodigal.^80^ To analyze the genomic features and calculate various sizes such as the size of CDS, tRNA, and IGS, we employed a custom python script and used Prokka annotated FASTA and GFF files (available on GitHub repository). CDS and tRNA sizes were extracted from the GFF file by identifying feature types and calculating sizes based on start and end positions. IGS, defined as the segments of the genome between annotated CDS features, were calculated as the difference between the start position of the current CDS feature and the end position of the previous feature minus one.

To assess the deviation of IGS from CDS GC content, we extracted nucleotide sequences for CDS and IGS from FASTA files based on the coordinates in GFF files. We categorized IGS regions into those shorter than 300 base pairs, representing the typical size of bacterial IGS, and those longer than 300 base pairs, contributing to the low coding densities in *Richelia* genomes. The precise calculation procedure is available at GitHub repository.

### Transposable elements analysis

To identify transposases, genomes were compared to a database of insertion sequences with a two-step blastx process, using Transposeek2 [2024,https://github.com/Omnistudent/transposeek2]. The Transposeek2 python script compiles genomic footprints of regions with transposases, then divides these footprints into identified insertion sequences using the highest score. The amino acid database of transposases was originally downloaded from ISfinder [2016,https://isfinder.biotoul.fr/about.php].

### Pseudogenes identification

Pseudogenes were identified and annotated using Pseudofinder^81^ with default settings and by using NCBI nr database. Detailed descriptions of files produced by pseudofiner (v1.1.0) can be found https://github.com/filip-husnik/pseudofinder/wiki/5.-Commands, and all corresponding files from this project are available in the GitHub repository. Using the GFF files produced by Pseudofinder, we calculated the numbers and lengths of pseudogenes in each genome. Furthermore, we divided pseudogenes into three categories based on the provided justification for pseudogenization: run-on, truncated, predicted fragmentation, and blast hits in intergenic spacers. Pseudogenes annotated as transposases in the BlastP and BlastX files produced by Pseudofinder were excluded from GFF files based on their identifiers and therefore from the overall analysis. Detailed calculations of the above are represented in the python script available on GitHub.

### Pangenome analyses

Pangenome of the 4 draft *Richelia* genome assemblies and 10 recovered MAGs was computed using anvi’o^82^ standard pangenomics workflow (anvio V8) with some additions. ^82,83^ Briefly, we ran mOTUpan (v0.3.2) inside anvi’o with *anvi-script-compute-bayesian-pan-core* to computationally estimate whether gene clusters belong to the core or accessory genomes.^83^ This classification was then used to visualize the core genome within the pangenome (Figure 3). Briefly, the pangenome workflow consists of 1) generating contigs database out of genome FASTA files with program *anvi-gen-contigs-database* 2) generating genome storage database with program *anvi-gen-genomes-storage* 3) generate anvi’o pangenome database with program *anvi-pan-genome* 4) visualize pangenome using program *anvi-display-pan*. The detailed description of what program *anvio-pan-genom*e does can be found in Delmont and Eren, 2018.^83^ Briefly, it begins by calculating amino acid sequence similarities using blastP^85^, filters out weak hits, and then employs the MCL algorithm^86^ to identify gene clusters. The gene clusters, as described previously^83^, represent sequences of one or more predicted open reading frames grouped together based on their homology at the translated DNA sequence level. Subsequently, it computes the distribution of these gene clusters across genomes, conducts hierarchical clustering analyses for both gene clusters and genomes, and finally generates an anvi’o pan database which was used to visualize *Richelia* pangenome.

### Annotations

Genomes were initially annotated with Prokka and further annotated using EGG-NOG mapper (v2.1.9)^87,88^ with the reference database v5 and BLASTkoal (v 3.0).^89^ Prokka and emapper annotations were imported into pangenome as detailed in GitHub repository. Additionally, we performed functional annotations inside of anvio’o with COG annotations using the *anvi-run-ncbi-cogs* program with the –sensitive flag (runs sensitive version of DIAMOND (v2.1.8)^90^ and the 2020 COG20 database.^91,92^ KEGG/KOfam (v4)^93,94^, annotations were also added to each genome database file, as well as hmm-hits (v3.3.1).^86^ Summary file produced by anvio is available at repository. All our functional and metabolic analysis were based on annotations produced by emapper and KEGG database. All annotations discussed in this manuscript were further manually confirmed by inspecting their conserved domains using the NCBI conserved domain batch search.^95^

### KO annotations outside of the pangenomic pipeline

In order to compare the genomes of *Richelia* with other symbiotic cyanobacteria associated with diatoms and to avoid gene duplication, we downloaded amino acid FASTA files for *EtSB* (accession number GCA_000829235.1), *nitroplast* (*UCYN-A*, accession number GCA_020885515.1), and *N. azollae* (GCA_000196515.1). We annotated these genomes, along with all *Richelia* genomes, using the online version of BlastKOALA (v 3.0).^89^ Our functional analysis was based on the KEGG annotations generated from this process. Since many of the modules in the KEGG database are based on *E. coli* or other model bacteria, we took extra care when analyzing our pathways. Specifically, when we encountered missing genes, we did not immediately conclude that a pathway was incomplete. Instead, we only classified a pathway as incomplete after comparing them to genomes of well-studied cyanobacteria, such as *Synechocystis* sp. PCC 6803 (https://www.genome.jp/kegg-bin/show_organism?menu_type=genome_info&org=syn), *Nostoc* sp. PCC 7120 (*Anabaena* sp. PCC 7120, https://www.genome.jp/kegg-bin/show_organism?org=ana) and *Rivularia* sp. PCC 7116 (https://www.genome.jp/kegg-bin/show_organism?org=riv). While accounting for genome incompleteness, we considered a gene or function as truly absent in a group only if it is missing from all genomes within that group, and as present only if it appears in 90 % of genomes within that group. We also acknowledge the possibility that some functions may not be annotated, or that some genes for alternative pathways may have not yet been discovered or characterized.

### Computing average nucleotide identity

Similarity of genomes in the pangenome was calculated in anvi’o with anvi-compute-genome-similarity using PyANI (v0.2.13.1).^96^ This program calculated average nucleotide identity (ANI) which was used in for visualization together with the pangenome (Figure 4).

## Acknowledgements

The computations and data handling were enabled by resources in project NAISS 2023/23-493 and NAISS 2023/22-993 provided by the National Academic Infrastructure for Supercomputing in Sweden (NAISS) at UPPMAX, funded by the Swedish Research Council through grant agreement no. 2022-06725. Vesna Grujcic was supported by KAW Prolongation grant to RAF. Maliheh Mehrshad was supported by a grant from the Swedish Research Council for sustainable development, FORMAS (Grant no. 2021-00546).

## Author contributions

V.G., M.M. and R.A.F conceptualized the study. V.G, M.M., and D.L. performed bioinformatic analyses and developed figures. TVS analyzed insertion sequences. V.G., M.M., and R.A.F. interpreted the data and wrote the manuscript. All authors read and approved the manuscript.

## Supplementary Figures

**Supplementary figure 1.**
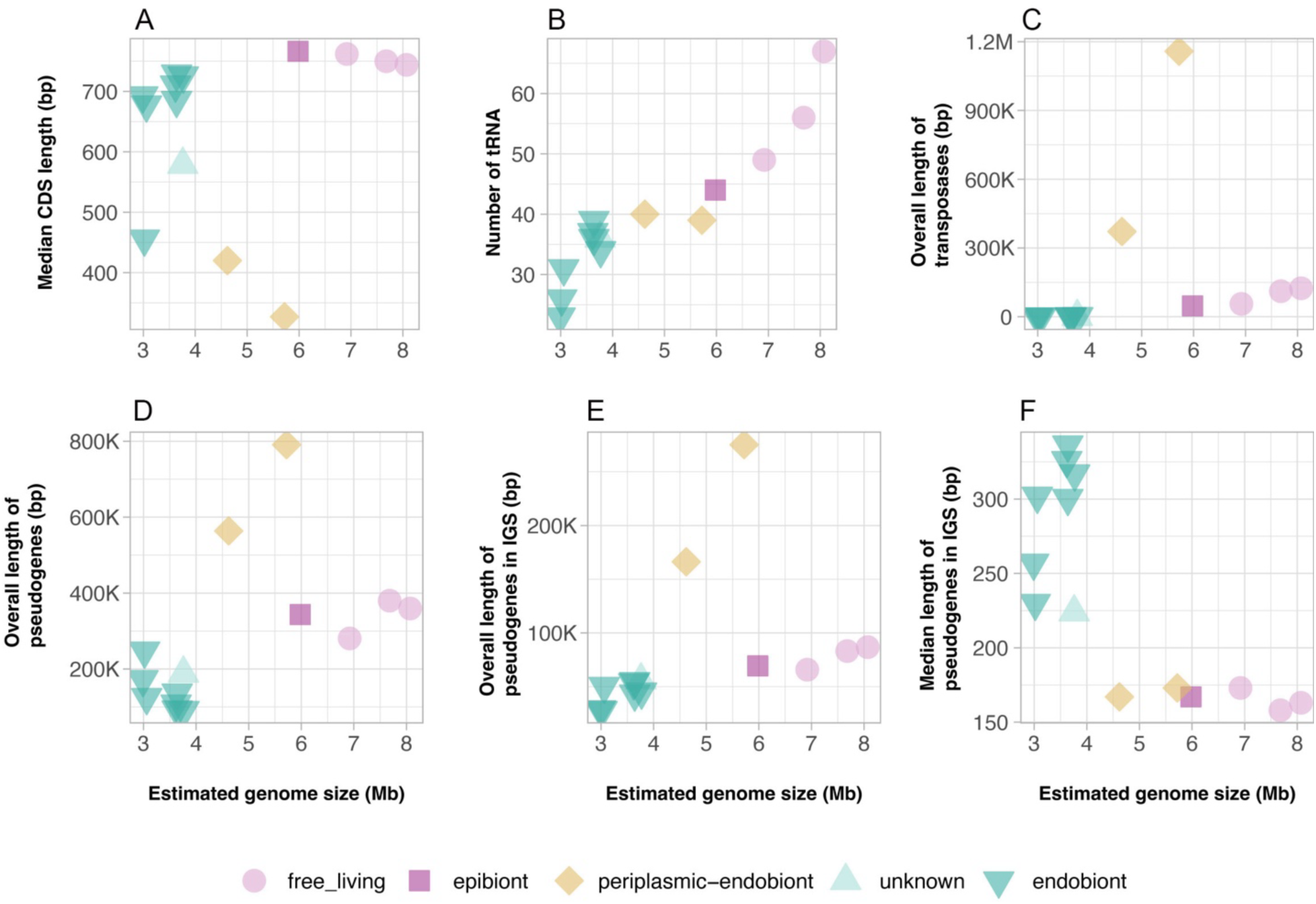
Genomic features of the *Richelia* spp. and eMAGs and reference genomes in relation to genome size. Each panel (a-g) represents a different genomic characteristic plotted against genome size. Different shapes and colors represent the categories: the epibiont and related eMAGs (diamonds), periplasmic-endobionts (triangles), and endobionts (squares).

**Supplementary figure 2.**
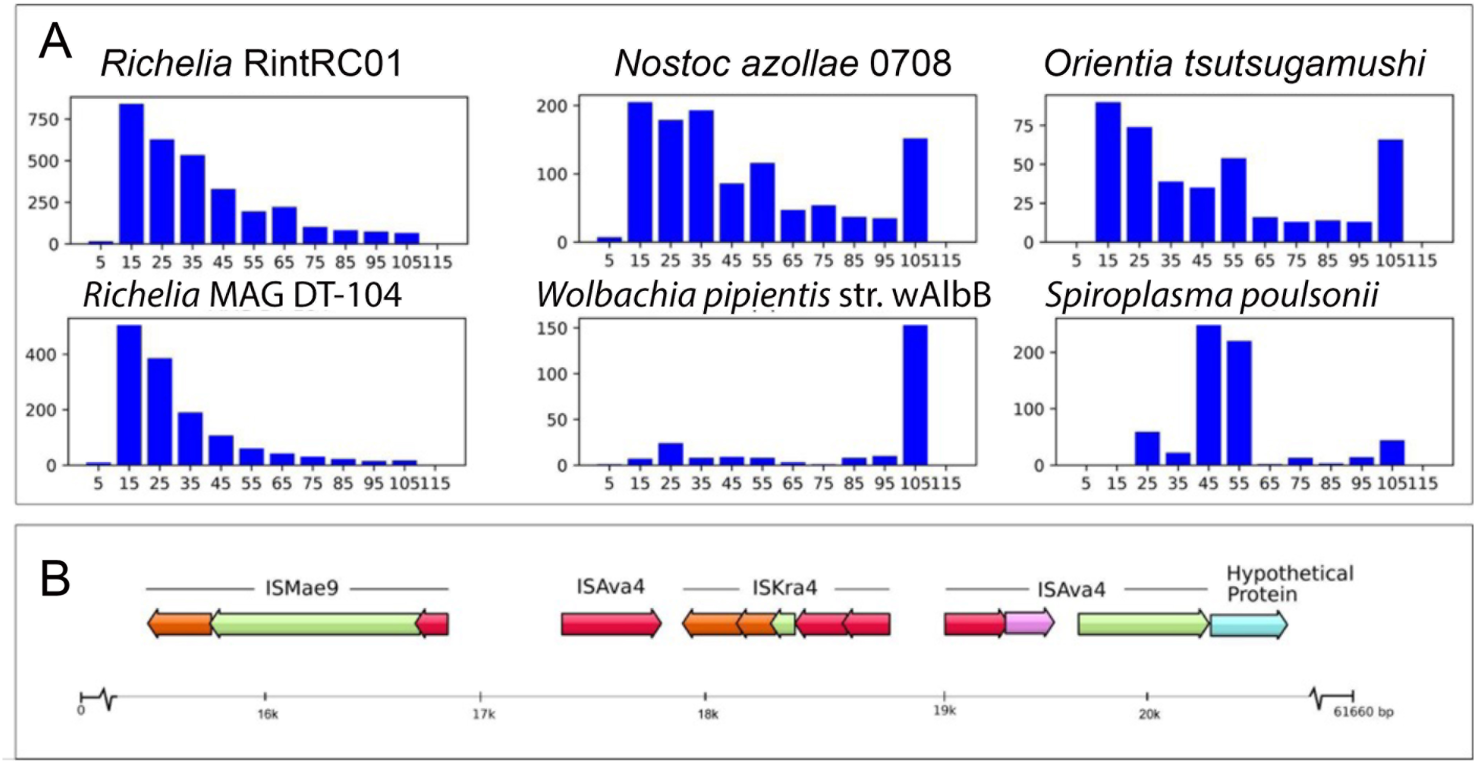
Summary of transposase (TE) analyses. a) Comparison of length distributions for the identified TEs in the periplasmic endobiont *Richelia* RintRC01 and different obligate symbiotic bacteria known to possess high numbers of TEs. X axis: length in percent of full TE length. Y axis: number of identified TEs with length ≤X%. b) Annotations for a TE rich part of contig650 from the genome of RintRC01, with annotations showing most similar TE, as named by ISFinder. Colors indicate: Turquoise - Prokka annotated CDS. Other colors represent what part of a TE a region originates from: red - start, purple - middle, orange - end, green - part unassigned.

**Supplementary figure 3.**
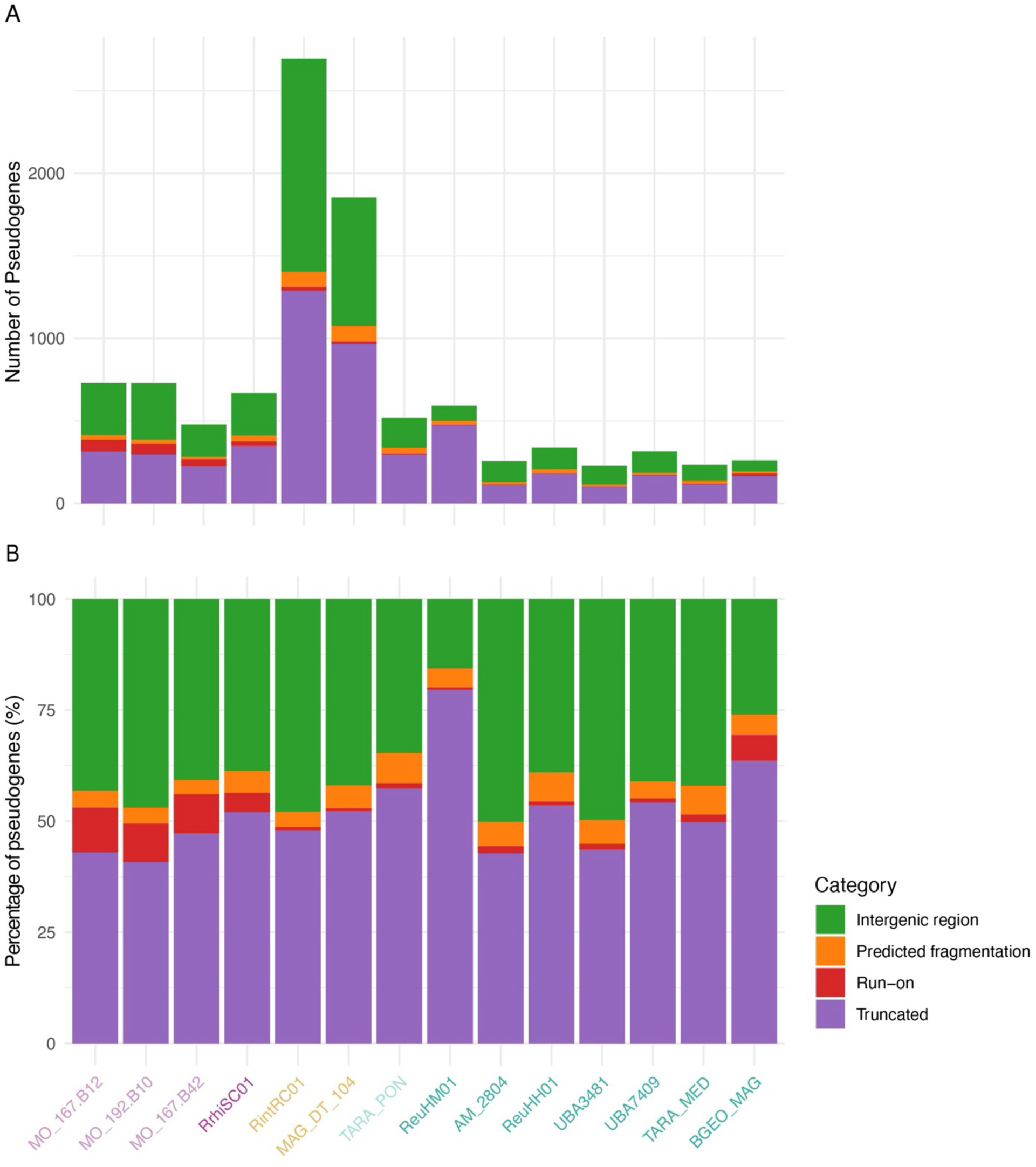
Pseudogene categorization generated by pseudofinder for Richelia spp. and eMAGs. a) The total count of pseudogenes categories per genome b) the percentage of pseudogene categories per each genome. Genomes (X-axis) are color-coded according to their cellular location: free-living (light purple), epibiont/facultative symbiont (purple), periplasmic or partial endobiont (yellow), endobionts (green) and unknown (light green).

**Supplementary figure 4.**
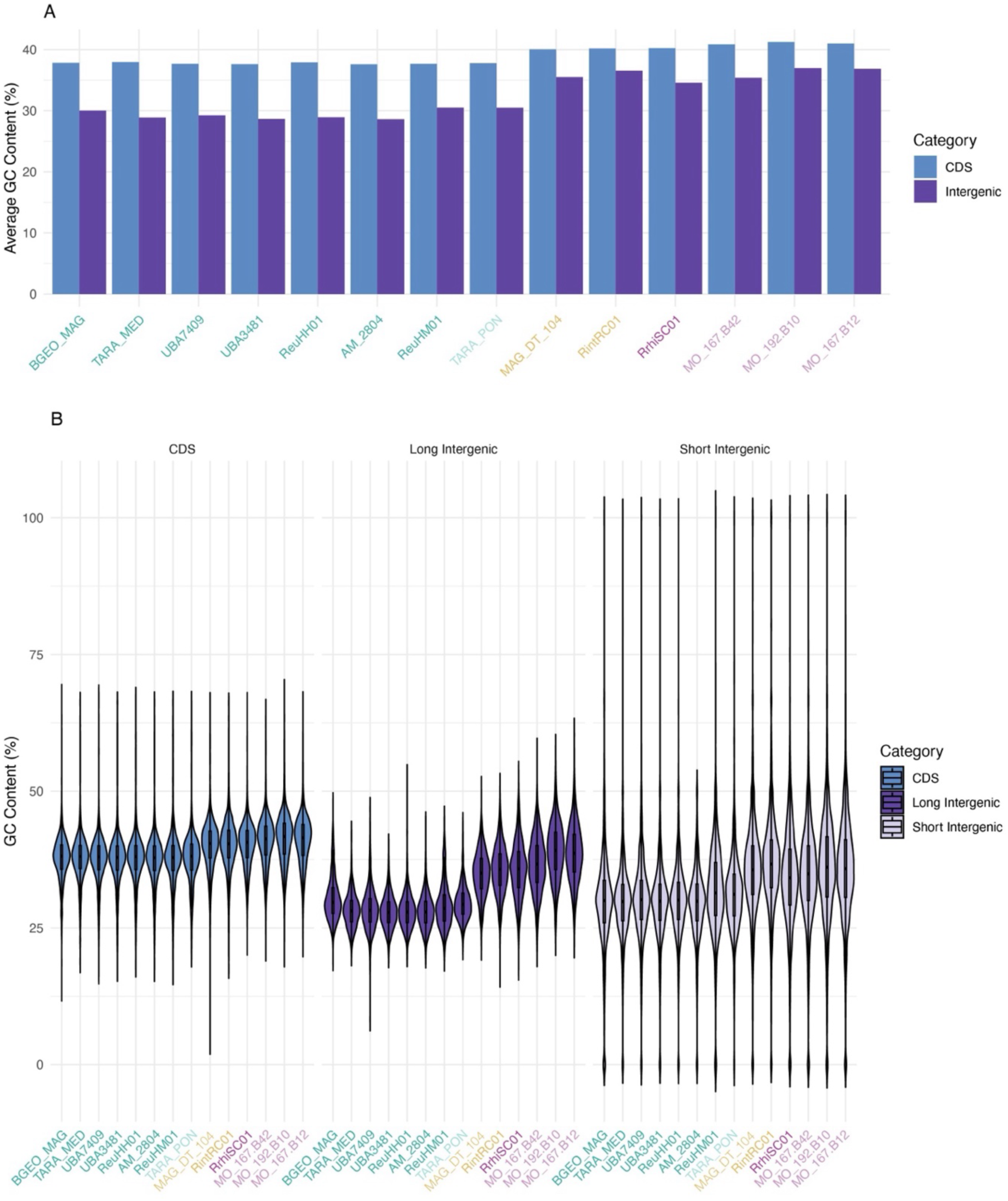
GC content distribution across coding sequence and intergenic spacers of the *Richelia* spp and eMAGs. a) Bar plots representing average GC content (%) across coding sequences (CDS) and intergenic regions for all the analyzed genomes. b) Violin plots showing detailed distribution of GC content in CDS, short intergenic regions (<300) and long intergenic regions (>300). Genomes are color-coded according to their cellular location: free-living (light purple), epibiont/facultative symbiont (purple), periplasmic or partial endobiont (yellow), endobionts (green) and unknown (light green).

**Supplementary figure 5.**
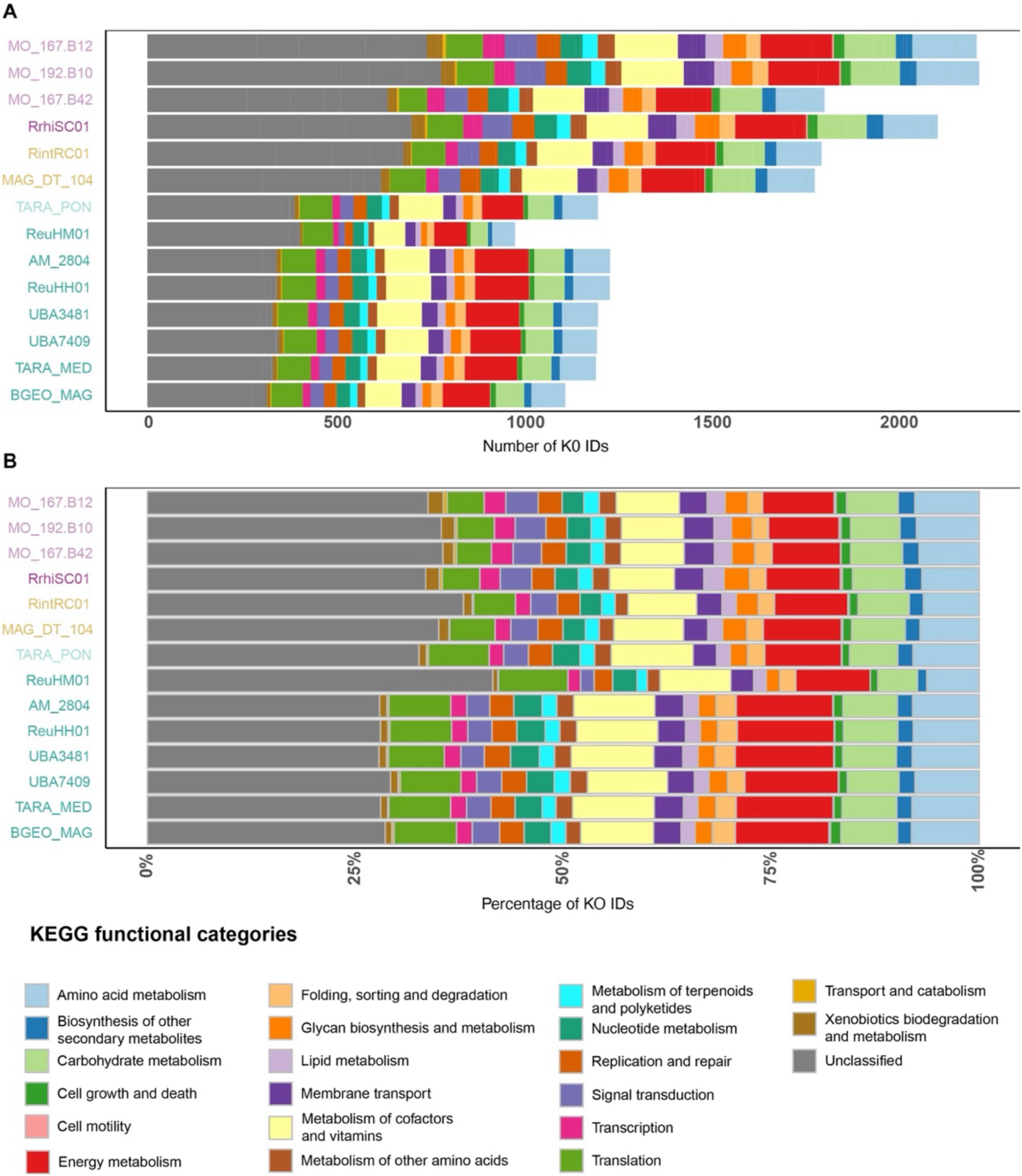
Distribution of gene clusters across KEGG functional categories in *Richelia* and eMAGs. a) An overview of the functional diversity across different genomes is shown as the total number of gene clusters assigned to each KEGG functional category per genome, b) The percentage of gene clusters within each KEGG functional category per genome illustrates the relative importance of different functional categories in each genome. Genomes are color-coded according to their cellular location: free-living (light purple), epibiont/facultative symbiont (purple), periplasmic or partial endobiont (yellow), endobionts (green) and unknown (light green).

**Supplementary figure 6.**
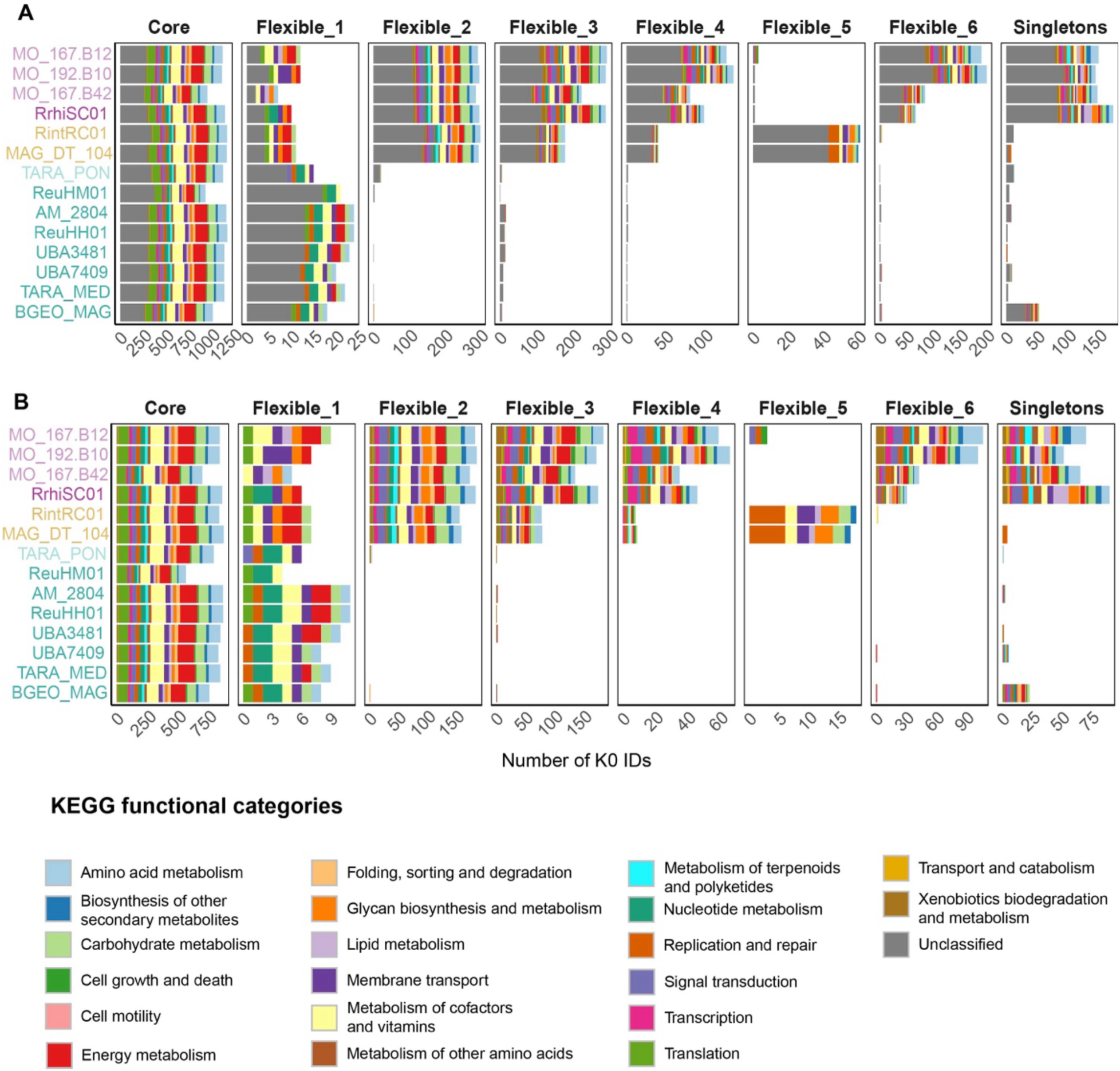
Distribution of gene clusters across KEGG functional categories in *Richelia* and eMAGs. a) The number of gene clusters assigned to each KEGG functional category across different pangenome sections: core, flexible (1-6), and singletons. Unclassified gene clusters are also included. b) The number of gene clusters assigned to each KEGG functional category in the core, flexible (1-6), and singleton fractions of the genomes, with unclassified gene clusters removed to enhance the visibility of the other functional categories. Genomes are color-coded according to their cellular location: free-living (light purple), epibiont/facultative symbiont (purple), periplasmic or partial endobiont (yellow), endobionts (green) and unknown (light green).

**Supplementary figure 8.**
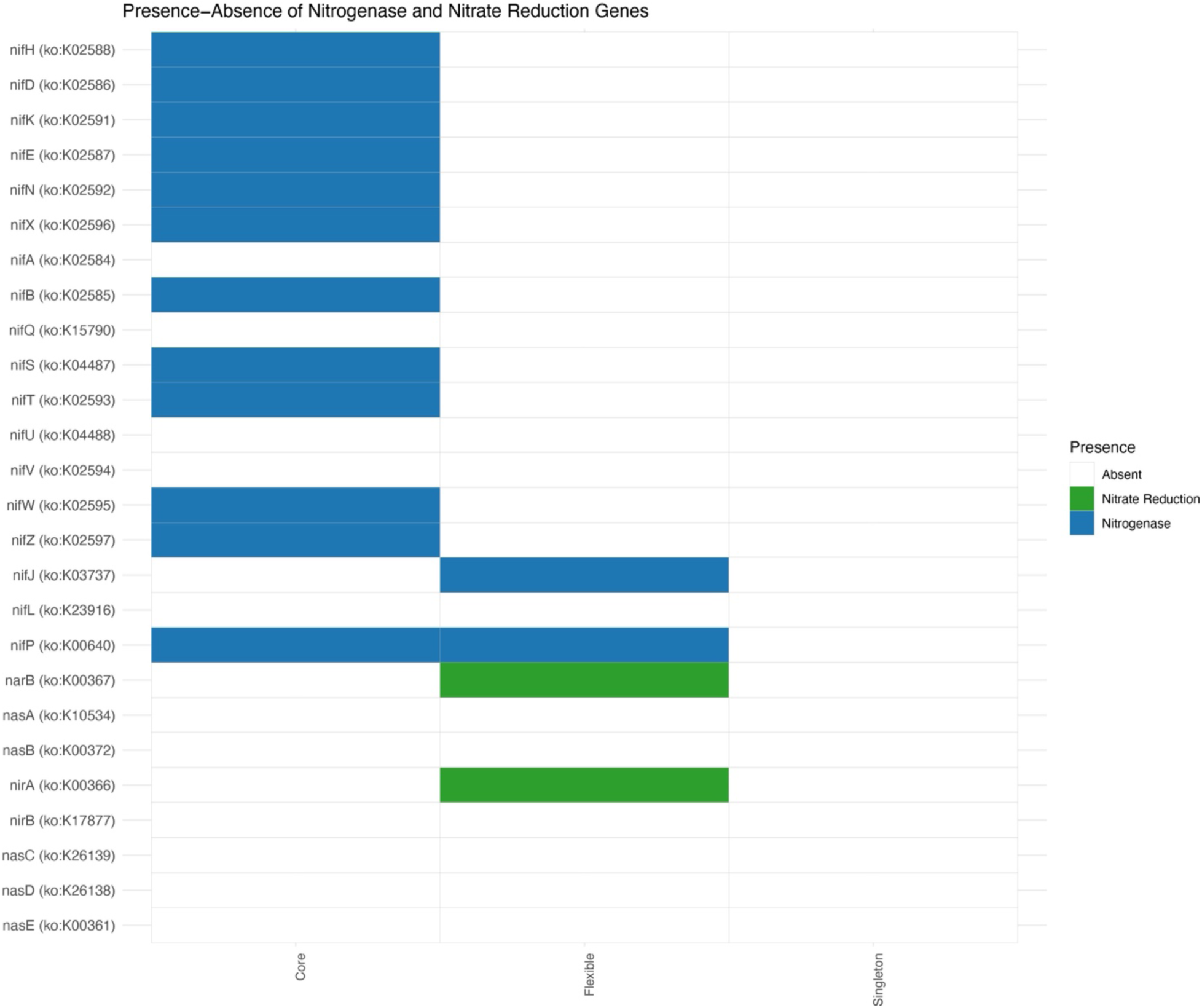
Presence-Absence of genes that encode for nitrogen fixation and nitrate assimilation across the pangenome of *Richelia*. The figure illustrates the presence (green) and absence (white) of genes in different sections of the *Richelia* pangenome: core, flexible, and singletons. Each row represents a specific gene (identified by its KEGG Orthology (KO) number), with columns showing the distribution of these genes within the pangenome bins - core, flexible, singleton.

**Supplementary figure 8.**
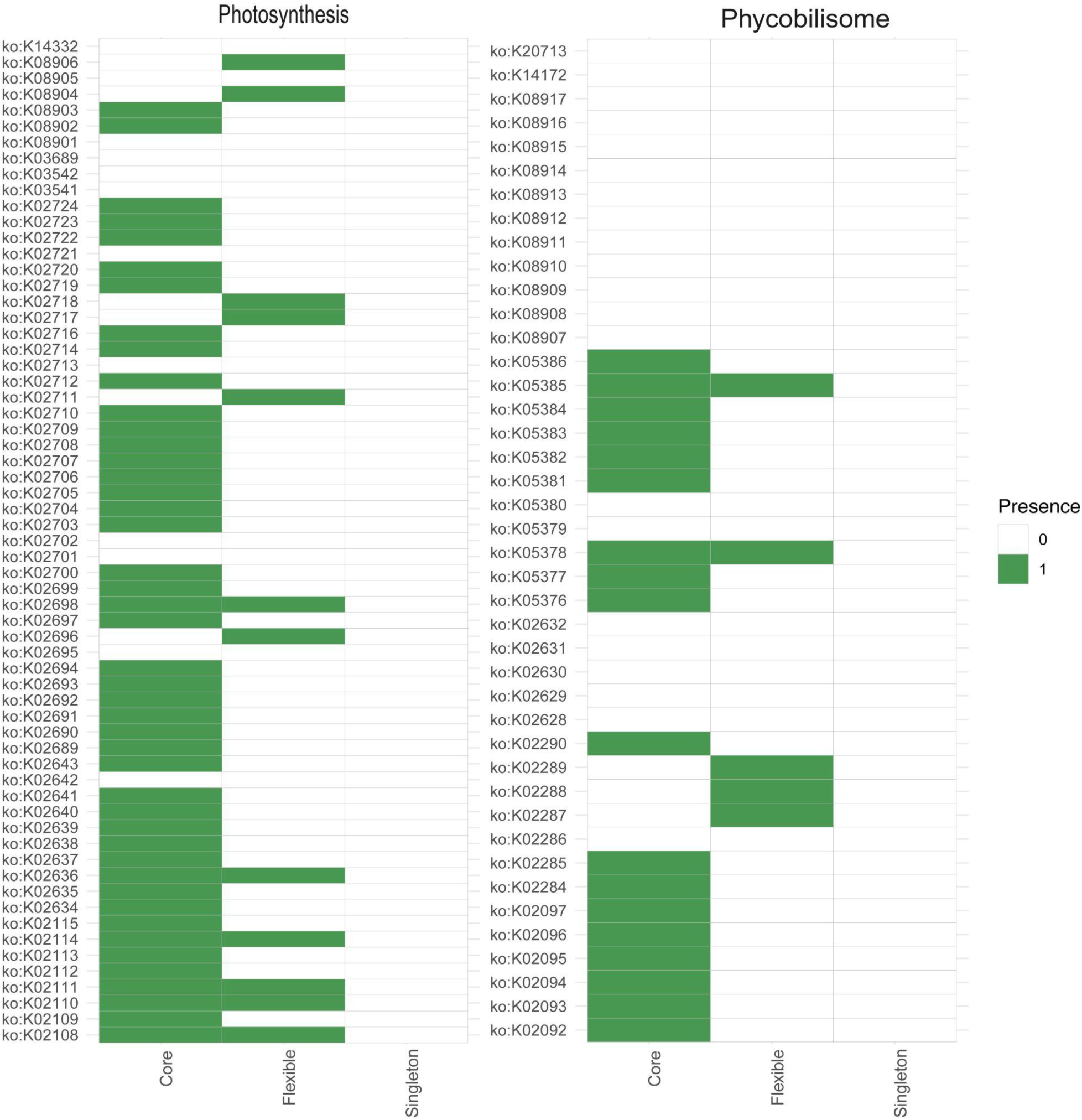
Presence-Absence of genes for photosynthesis and phycobilisomes in the pangenome of *Richelia*. The heatmap illustrates the presence (green) and absence (white) of genes associated with the KEGG photosynthesis and phycobilisome modules in the *Richelia* pangenome: core, flexible, and singletons. Each row represents a specific gene (identified by its KEGG Orthology (KO) number), with columns showing the distribution of these genes within the pangenome bins - core, flexible, singleton.

**Supplementary figure 9.**
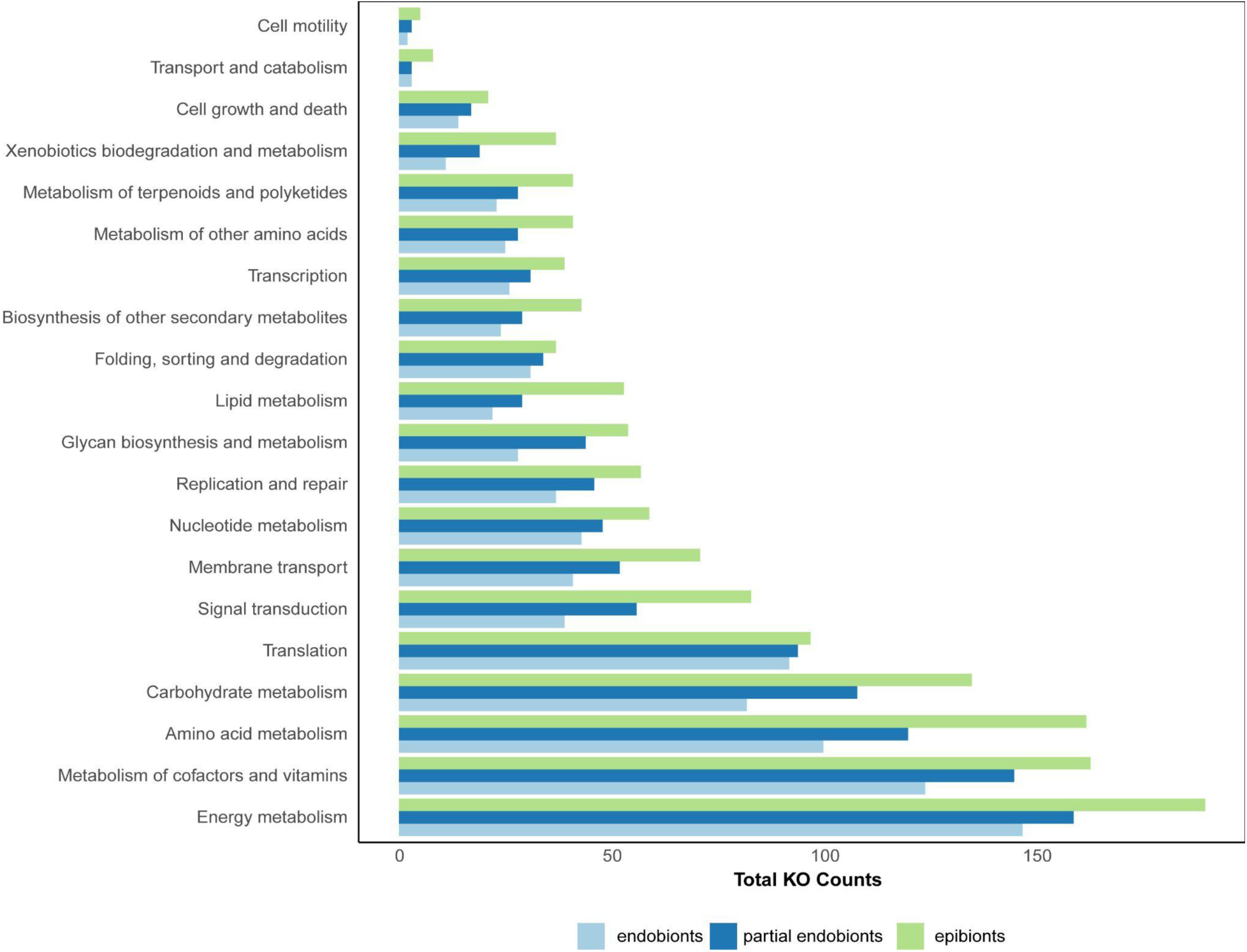
Total KEGG Orthology (KO) counts in the various functional categories of endobionts, periplasmic endobionts, and the epibiont. A comparison of the total number of KO associated with key metabolic and functional categories across different symbiotic categories: endobionts (light blue), partial endobionts (dark blue), and the epibiont (light green).

**Supplementary figure 10.**
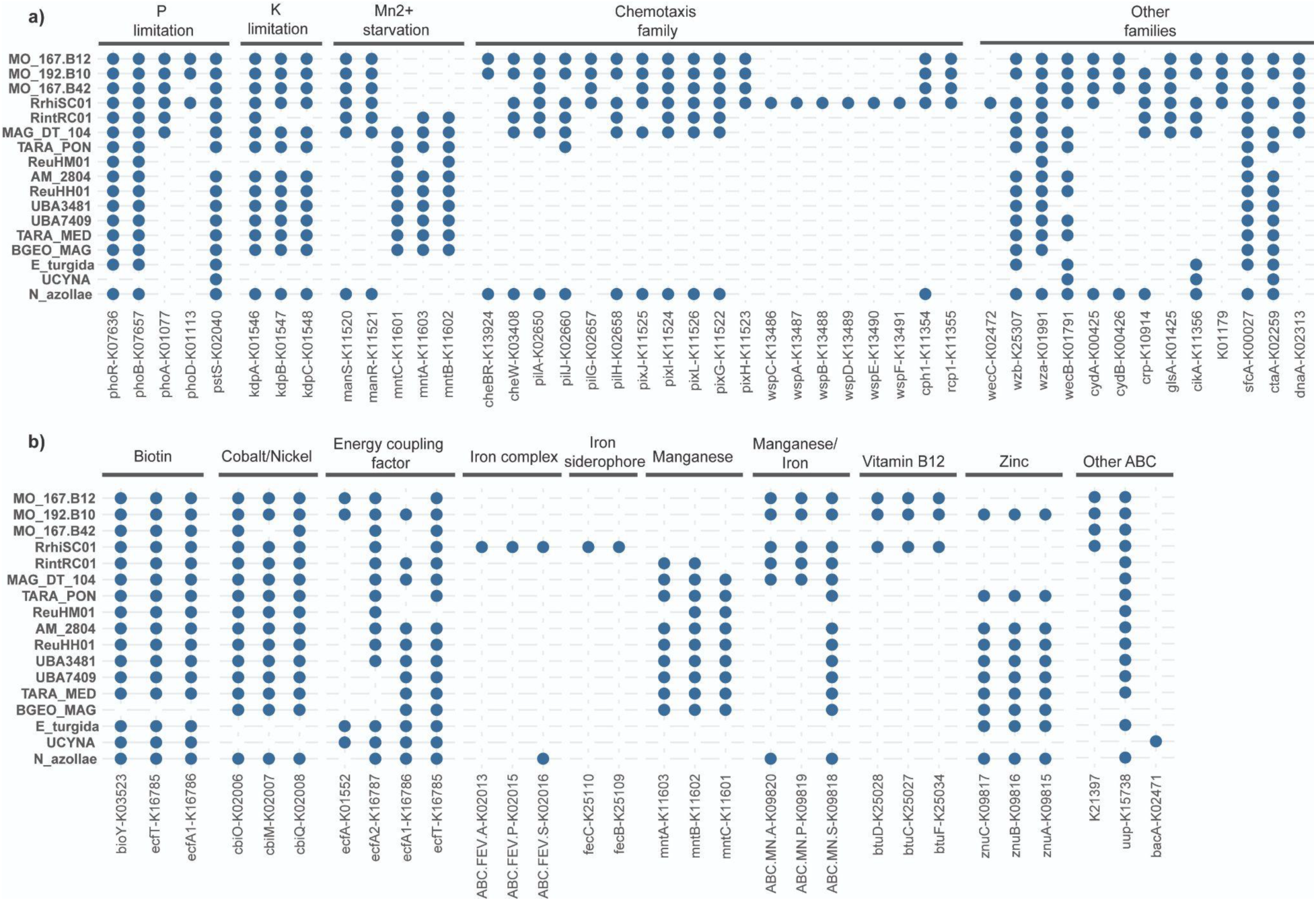
Presence-Absence of genes that encode the two-component secretion systems (a) and some specific transporters in the *Richelia* genomes compared with other symbiotic cyanobacteria and UCYN-A/Nitroplast. **(b)**. K0 number identifiers together with corresponding gene names are shown as column names. *Richelia* genomes, the free-living eMAGs, and three other genomes belonging to other symbiotic cyanobacteria and UCYN-A/Nitroplast are represented by row names. Gene presence is marked as a blue circle and absence as an empty space.

## Supplementary tables

**Supplementary table 1. Genome information and statistics of all the *Richelia* genomes and eMAGs analyzed in this study.** A detailed overview of genomic statistics across various *Richelia spp.* strains and eMAGs, and includes accession numbers and taxonomy classification. Genomes in the table are ordered, as elsewhere in the manuscript, by their cellular location: free-living, epibiont, periplasmic endobionts, or endobionts. Genome statistics presented in this table were used to produce Figure 2 and Supplementary figure 1.

**Supplementary table 2. Transposable element (TE) statistics and associated genome metrics for periplasmic endobionts and other obligate symbionts and parasites known to have high numers of TEs.** A summary of key genomic features, including TE content as a percentage (%) of the genome, GC content (%), genome size (Mbp), total TE footprint (bp), median transposase length (bp), and the number of contigs in each genome assembly.

**Supplementary table 3. Numbers of pseudogenes belonging to different categories per genome.** A summary of the distribution of pseudogenes across various categories within the genomes analyzed. The numbers were extracted from the output produced by pseudofinder program. The pseudogenes are classified into different categories based on the reasons for their formation, which include intergenic regions, predicted fragmentation, run-on transcription, and truncation. Each genome’s count of pseudogenes in these categories is provided, together with the overall length (bp) and median length (bp).

**Supplementary table 4. Distribution of gene clusters across different bins in various genomes.** The distribution of gene clusters across core, flexible (accumulated flexible 1-6), and singleton bins for the different *Richelia* genomes and eMAGs. It includes the total number of gene clusters per genome, the number of clusters within each bin, and the corresponding percentage that each bin represents in the genome.

